# Enhanced short-wavelength sensitivity in the blue-tongued skink, *Tiliqua rugosa*

**DOI:** 10.1101/2022.03.25.485754

**Authors:** Nicolas Nagloo, Jessica K. Mountford, Ben J. Gundry, Nathan S. Hart, Wayne I. L. Davies, Shaun P. Collin, Jan M. Hemmi

## Abstract

The complex visually mediated behaviors of diurnal lizards are enabled by a retina typically containing five types of opsins with the potential for tetrachromatic color vision. Despite lizards using a wide range of color signals, the limited variation in photoreceptor spectral sensitivities across lizards suggests only weak selection for species-specific, spectral tuning of photoreceptors. Some species, however, have enhanced short wavelength sensitivity, which likely helps with the detection of signals rich in ultraviolet and short wavelengths. In this study, we examined the visual system of *Tiliqua rugosa*, which has a UV/blue tongue, to determine the spectral sensitivity of the eye and to gain insight into this species’ visual ecology. Electroretinograms coupled with spectral stimulation showed peak sensitivity at 560 nm with high similarity to other lizards at wavelengths greater than 530 nm. However, at shorter wavelengths, sensitivity is enhanced leading to a spectral sensitivity curve that is 28 nm broader (full width at half height) than other lizards studied so far. The width of the curve is partially explained by a population of photoreceptors that respond more strongly to low temporal frequencies with possible peaks in sensitivity between 460 and 470 nm suggesting that they are SWS2 photoreceptors. The lack of a peak in sensitivity at 360 nm at low temporal frequencies and under a monochromatic light that suppresses the response of LWS photoreceptors, suggests that the SWS1 photoreceptors are red-shifted. In addition, the yellow and green oil droplets that are common in other diurnal lizards appear to be missing and instead, only transparent and pale-yellow oil droplets are present. LWS photoreceptors are likely paired with pale-yellow oil droplets to produce LWS photoreceptors with wider spectral sensitivity curves than in other lizards. Opsin sequencing reveals *SWS1, SWS2, RH1, RH2* and *LWS* opsin genes that are very similar to the visual opsins detected in the green anole, *Anolis carolinensis*, suggesting there is little change in the spectral sensitivity of photoreceptors compared to other diurnal lizards. Since we only obtained a partial sequence of the SWS1 opsin, we were unable to determine whether amino acid substitution at tuning sites could have played a role in red-shifting the SWS1 photoreceptor spectral sensitivity. Photoreceptor densities are typically higher in central and ventral retinal regions than in the dorsal retina suggesting that higher spatial sampling is necessary at eye level and above the animal than on the ground. However, the SWS1 photoreceptors do not follow this pattern potentially due to their low abundance making them less relevant to high acuity visual tasks. Our findings demonstrate that there are possibly multiple mechanisms acting synergistically in the visual system of *T. rugosa* to enhance short wavelength sensitivity between 360 and 530 nm. While it is tempting to suggest that this is an adaptation to facilitate the detection of the blue tongues of conspecifics, additional experiments are necessary to determine its ecological relevance.

**Summary statement:** Color vision and the distribution of photoreceptor subtypes in *T. rugosa*

## 1 Introduction

There are approximately 8000 reptilian species that occupy a range of aquatic and terrestrial environments (Hickman et al., 2008). Amongst them, lizards are the most ecologically diverse group with representatives occupying a range of terrestrial, fossorial, aquatic, arboreal and aerial habitats (Hickman et al., 2008). In many species examined thus far, adaptive radiation of the visual system appears to mirror the ecological diversity of lizards (Walls, 1942), where vision plays a crucial role in predation and conspecific communication (Kirmse et al., 1994; Stapley and Whiting, 2006). However, previous studies have mainly focused on well-known subgroups such as geckoes, iguanids, and chameleons (Crescitelli, 1977; Govardovskii et al., 1984; Loew et al., 2002). As such, there is a clear gap in our understanding of visual ecology for many other common and ecologically important subgroups of reptiles.

During important intra-specific interactions, such as courtship displays and male-to-male fitness signaling, lizards use a variety of colorful conspecific signals which often extend beyond the visible spectrum and into the ultraviolet. This has led visual ecologists to investigate whether lizard visual systems are spectrally tuned to facilitate the detection of these signals (Fleishman et al., 1997; Loew et al., 2002). Previous experiments have used microspectrophotometry (MSP) to obtain the spectral sensitivity of specific photopigments, photoreceptor types and oil droplets (Barbour et al., 2002; Govardovskii et al., 1984; Katti et al., 2019; Kawamura and Yokoyama, 1998; Takenaka and Yokoyama, 2007) or electroretinograms (ERG) to measure the spectral sensitivity of the whole eye (Arden and Tansley, 1962; Ellingson et al., 1995; Fleishman et al., 1997; Forbes et al., 1960; Hamasaki, 1968).

Data generated using MSP from 17 species of Caribbean anole lizards reveals the presence of four spectrally distinct photoreceptor types with little variation in peak sensitivities across species, despite the great variety of dewlap colors involved in conspecific signaling. The wavelength of peak sensitivity (λ_max_) for the four photoreceptor types are at 365 nm (ultraviolet-sensitive, UVS), 456 nm (short-wavelength-sensitive, SWS), 494 nm (middle-wavelength-sensitive, MWS), and 564 nm (long-wavelength-sensitive, LWS) (Loew et al., 2002). The λ_max_ of these photoreceptor types seem to also be broadly conserved in geckoes, iguanids, skinks and lacertids (Martin et al., 2015) with only small deviations. The four spectrally-distinct photoreceptor types typically found in diurnal lizards are generally the result of the expression of four cone opsin genes, namely a long-wavelength-sensitive (*LWS*) opsin, a rhodopsin-like 2 (*RH2*) opsin, short-wavelength-sensitive 2 (*SWS2*) and short-wavelength-sensitive 1 (*SWS1*) opsins, and a single rod (*RH1*) opsin gene (Katti et al., 2019; Kawamura and Yokoyama, 1997; Martin et al., 2015; Yewers et al., 2015). Unlike most vertebrates that possess rod photoreceptors that express *RH1*, the rod opsin gene in some species of lizards is expressed in single, cone-like photoreceptors, an observation that is widespread across diurnal lizards, i.e. in *Chamaeleo chamaeleon, Anolis carolinensis* and *Tiliqua rugosa* (Bennis et al., 2005; McDevitt et al., 1993; New et al., 2012). At the amino acid level, visual opsins show high sequence identity within each opsin class across lizard species. These opsins are bound to a retinal-based chromophore which can shift the spectral peak and broaden the bandwidth of the individual photopigment class. While most terrestrial vertebrates, including diurnal lizards, are known to use an A_1_-based chromophore (i.e. 11-*cis* retinal), mixtures of both A_1_ and A_2_ (3,4-didehydroretinal) chromophores have been observed in *Podarcis sicula, Chameleo dilepis* and *Fucifer pardalis* (Bowmaker et al., 2005; Provencio et al., 1992). Pure A_2_ chromophore is found only in *A. carolinensis* and *Zootoca vivipara* (Kawamura and Yokoyama, 1993; Loew et al., 2002; Martin et al., 2015). The combination of A_1_ and/or A_2_ chromophores bound to visual opsins is therefore an important mechanism of adaptive spectral tuning (Corbo, 2021; Hárosi, 1994; Whitmore and Bowmaker, 1989).

ERGs of anole lizards (*A. gundlachi, A. cristatellus, A. krugi, A. pulchellus, A. stratulus, A. evermanni* and *A. sagrei*), horned toads (*Phrynosoma spp*.), spiny lizards (*Scleroporus spp*.) and geckoes (*Gonatodes albogularis*), suggest that the spectral sensitivity of the whole eye, is as conserved as the spectral sensitivity of photoreceptor types (Arden and Tansley, 1962; Ellingson et al., 1995; Fleishman et al., 1997; Forbes et al., 1960). The typical spectral sensitivity curve of the diurnal lizard eye is characterized by a broad shoulder of high sensitivity between 530 nm and 590 nm and a secondary (less sensitive) peak at approximately 360 nm (Fleishman et al., 1997). Fleishman et al. (1997) have hypothesized that the convergence of these spectral sensitivity curves may be driven by the common need to detect objects against green vegetation in the background, which has a peak reflectance at 550 nm (Fleishman et al., 1997). Despite this pattern of conserved spectral sensitivity, a more recent study has revealed enhanced sensitivity at 360 nm (ultraviolet) in the retina of the cordylid lizard, *Platysaurus broadleyi* (Fleishman et al., 2011), which correlates with an increased abundance of UVS receptors. While color discrimination models have revealed that this enhanced sensitivity would facilitate the detection of conspecific signals rich in UV, it remains difficult to determine whether the spectral properties of conspecific signals can shape the spectral sensitivity of diurnal lizards. Additional studies examining the spectral sensitivity of the whole eye across lizard taxa, which make use of conspicuous visual signals, are necessary to untangle the multiple factors which drive spectral sensitivity and signal detection in diurnal lizards.

Sleepy lizards, *Tiliqua rugosa*, are a species of blue-tongued lizard where males are highly aggressive towards each other with interactions that lead to scarring and scale loss (Murray and Bull, 2004). Blue-tongue displays are thought to play a role in male-to-male signals and could be used to avoid costly physical altercations during the mating season (Abramjan et al., 2015; Murray and Bull, 2004). Recent work in the common blue tongue lizard *T. scincoides* also suggests that the tongue might be a deterrent to predators by providing a sudden overwhelming flash of UV and blue light that would intimidate and startle predators (Badiane et al., 2018). These blue tongues are not limited to the *Tiliqua* genus but have also been observed in closely related large skinks with tongue color ranging from pale gray to dark blue with primary peak reflectance at ∼320 nm and a secondary peak at ∼460 nm (Abramjan et al., 2015). The presence of blue tongues across several genera and their conspicuous nature offers an opportunity to closely examine the phylogenetic and environmental factors which drive whole-eye spectral sensitivities in these lizards. Here, we propose to start with a thorough examination of the visual system of *T. rugosa* to build on prior knowledge of its retinal organization.

New et al. (2012), previously showed that the retina of the sleepy lizard contains only cones with ∼20% of the cone population expressing the RH1 opsin. Both single and double cones were observed with the presence of pale-yellow oil droplets reported in single cones and the principal member of the double cones (New et al., 2012). The density of cones and retinal ganglion cells both peak in the retinal center with densities reaching 76,000 cells/mm^2^ and 15,500 cells/mm^2^, respectively (New and Bull, 2011; New et al., 2012). Anatomical estimates of the visual acuity suggest the sleepy lizards have a spatial resolving power of 6.8 cycles/degree (New and Bull, 2011). The current study used ERGs to measure the spectral sensitivity of the eye, opsin sequencing to characterize the full complement of opsin genes expressed, and immunohistochemistry and design-based stereology to visualize and map the complement of photoreceptors. The findings are discussed in relation to the visual ecology and behavior of *T. rugosa* in comparison to other lizard species.

## 2 Materials and Methods

### 2.1 Animals

Seven adult *T. rugosa* were caught around the Perth metropolitan area (Western Australia, Australia) and housed on the campus of The University of Western Australia for less than one year. An ultraviolet-emitting light globe (Repti Glo UVB10 Compact 13W, Exo-Tera, Canada) provided animals with a 12:12 light:dark (LD) photoperiod. All captive lizards were mature adults with a body weight ranging from 450 g to 670 g, and fed on a diet of soft vegetables supplemented with crickets and vitamins. All experimental procedures were approved by the ethics committee of The University of Western Australia (AEC No: RA/3/100/1030). All animals used in this study were euthanized according to protocols outlined in the ethics application above, with an intravenous injection of sodium pentobarbital.

### 2.2 Transmission of the optical elements of the eye

The transmission of the lens and cornea in the eyes of *T. rugosa* was measured using frozen samples. While the transmission of the vitreous can be affected by freezing, the lens and cornea remain largely unchanged, especially at shorter wavelengths (Pérez i de Lanuza and Font, 2014). The lens and cornea were thawed and placed on a perforated plate underneath an inverted integrating sphere (FOIS-1, Ocean Optics, USA). A 600 µm fiber (P600-2-UV-Vis, Ocean Optics, USA) carrying light from a pulsed-xenon arc lamp (PX-2, Ocean Optics, USA) was aligned to the center of the sample against the bottom surface so that light transmitted through the sample entered directly into the integrating sphere. The light from the integrating sphere was collected by a second 600 µm fiber (P600-2-UV-Vis, Ocean Optics, USA) and carried to a spectrophotometer (USB2000+, Ocean Optics, USA) for spectral analysis using SpectraSuite software (Ocean Optics, USA). At the beginning of experimental measurements, light and dark references were taken to calibrate the spectrophotometer to ambient light conditions.

The spectral transmittance (330-800 nm) of the different retinal cone photoreceptor oil droplets was measured using a single-beam wavelength-scanning microspectrophotometer as described previously (Hart, 2004). Briefly, small pieces of retina were dissected out of a fresh eyecup and mounted between coverslips in a solution of 8% dextran in 0.1M phosphate buffer saline (pH 7.2). Oil droplet transmittance was measured against a reference measurement made through a patch of nearby retinal tissue to control for absorbance of retinal tissue. While we were able to reliably record from two types of oil droplets, the process was inherently difficult due to the small size and fragility of reptilian oil droplets. We are unlikely to have missed additional oil droplet types due to a size bias as colorless or pale oil droplets tend to be smaller than the more conspicuous green, orange, red or yellow oil droplets (Goldsmith et al., 1984).

### 2.3 Measuring spectral sensitivity using electroretinograms

Animals were anesthetized with a combination of ketamine (Ceva Ketamine Injection, Australia; dosage: 50 mg/kg) and medetomidine (Ilium Medetomidine Injection, Australia; dosage: 165 µg/kg) administered intramuscularly during all electrophysiological experiments. Proxymetacaine hydrochloride (Alcaine 0.5%, Alcon, USA) was applied to the corneal surface to provide additional local anesthesia. A platinum active electrode was placed at the corneal surface with conductive gel, while a silver/silver-chloride reference electrode was placed behind the head and the ground connected to the Faraday cage. A differential amplifier (DAM50, World Precision Instruments, USA) combined with a National Instruments data acquisition board (USB-6353, National Instruments, USA) were used to amplify and collect the compound neural responses of the eye.

Spectral sensitivity was measured between 650 and 350 nm at 20 nm intervals and then from 340-640 nm at 20 nm intervals following a similar protocol and equipment used in previous studies (Jessop et al., 2020; Ogawa et al., 2015). The data was then spliced into a single spectral sensitivity curve from 350-650 nm with interleaved measurements every 10 nm. The stimulus consisted of an on/off flickering light alternating between colored and white light and separated by dark intervals of equal duration produced by chopping the light path with a motorized wheel (Jacobs et al., 1996). Time intervals between white and colored flashes were used to calculate signal frequency, with standard measurements carried out at 10 Hz. Monochromatic light (15 nm full width at half maximum transmission) was produced by an automated monochromator (Polychrome V, Till Photonics, USA), while white light was produced by a Xenon arc lamp (HPX-2000, Ocean Optics, USA). The two light paths were combined using mirrors and a beam splitter and gathered into a 1.1 mm quartz optic fiber (Till Photonics, USA). The end of the optical fiber was attached to a UV-transmitting quartz lens, which was placed ∼1.5 cm away from the eye. The proximity of the output fiber to the eye ensured that the light diverged and diffusely stimulated the retina at the back of the eye with colored and white light stimulating the same retinal region. Irradiance of the colored and white lights were measured using a ILT1700 radiometer with an SED033 detector combined with a flat response filter and a diffuser to achieve a cosine response profile (International Light Technologies Inc., USA). White light output from the optical setup reached a maximum irradiance of 3.11×10^−03^ W/cm^2^. The white light intensity could be adjusted using neutral density (ND) filters ranging from 0.8-1.3 ND to allow for a stronger white light response when adaptation lights were used but was held constant throughout a full spectral measurement. The irradiance of the colored lights ranged from 1.76×10^−03^ W/cm^2^ to 7.25×10^−03^ W/cm^2^ for wavelengths ranging from 350 nm to 650 nm and were dynamically adjusted to produce the same response amplitude as white light. At each 10 nm interval from 350 nm to 650 nm, sensitivity was measured as the reciprocal of the number of photons needed for the monochromatic light to produce an equivalent electrical response to that produced by white light (Neitz et al., 1989).

To analyze the contribution of specific photoreceptor subpopulations to the overall response of the eye, spectral sensitivity was measured under a range of conditions designed to alter the contribution of different photoreceptor populations to the recorded signal. To isolate spectrally-distinct photoreceptor types, a second monochromatic light (adaptation light from a Polychrome V, Till photonics, USA) was superimposed on the stimulus to selectively reduce contrast to cell populations that are maximally sensitive to different regions of the spectrum. In addition, the flicker rate of the stimulus was adjusted from 3 Hz to 30 Hz by controlling the speed of the chopper wheel. This was expected to bias contributions from slow or fast photoreceptor populations to the overall ERG signal (Neitz et al., 1989). At the beginning of each experiment, animals were dark adapted for 1 h and a standard spectral sensitivity curve recorded. After each use of a bright adaptation light, the animal was dark adapted for at least a further 30 min before the next recording.

### 2.4 Statistical comparison of spectral sensitivities across temporal frequencies

For any given comparison, the ERG curves were normalized by linearly fitting them to the average of all curves being compared. The ratio of short-to long-wavelength (SW:LW) sensitivity was then calculated and used to compare ERG responses across temporal frequencies. In this case, the ratio was obtained by comparing the integral of the curve on either side of 530 nm. This is halfway between the peak sensitivity of MWS (494 nm) and LWS (564 nm) photoreceptors previously reported in anole lizards (Loew et al., 2002) and allows for the comparison of LWS photoreceptors to all other photoreceptor subtypes. Reciprocal transformation of ratio data was used to reduce skewness in raw data. A generalized linear mixed effect model was then used on the transformed data temporal frequency as fixed effect and with individual identity as a random effect. This was implemented using the *fitglme* function in Matlab 2021a (Mathworks, USA) with the following model definition: fitglme(data, SWratio∼temporal_frequency + (1|animal_id)).

### 2.5 Isolation and sequencing of opsin mRNA

Eyes used for opsin sequencing were dissected and preserved in RNAlater (Sigma, Australia) at 4°C immediately after euthanasia of the animal and following enucleation of the eye. Total RNA was extracted from homogenized retinal samples using TRIzol Reagent with the PureLink RNA Mini Kit (Thermo Fisher Scientific, Australia), following the steps outlined by the manufacturer. Complementary DNA (cDNA) was subsequently generated using 2 μg of total RNA and the miScript II Reverse Transcription (RT) Kit (Qiagen, Australia), according to the manufacturer’s instructions. The *LWS, SWS1, SWS2, RH2* and *RH1* genes were PCR amplified according to Davies et al. (2009) from cDNA using degenerate primers listed in **Supplementary Table S1** and as follows: DIAPLMF1, DIAPLMF2, DIAPLMR1 and DIAPLMR2 for the amplification of the *LWS* gene; DIAPS1F1, DIAPS1F2, DIAPS1R1 and DIAPS1R2 for the *SWS1*; DIAPS2F1, DIAPS2F2, DIAPS2R1 and DIAPS2R2 for *SWS2*; DIAPPR2F1, DIPAPR2F2, DIAPR2R1 and DIAPR1R2 for *RH2*; and DIAPPR1F1, DIPAPR1F2, DIAPR1R1 and DIAPR1R2 for the amplification of the rod (*RH1*) gene (Davies et al., 2009; Hart et al., 2016; Knott et al., 2013). To confirm that a full complement of visual opsin genes was detected, a series of PCR experiments using AOAS primers (**Table S1**) were also performed. All PCR protocols and conditions are outlined in Davies et al. (2009): briefly, a first-round PCR was carried out using 200 ng of template cDNA with My Taq DNA polymerase (Bioline Alexandria, NSW, Australia), initial denaturation at 95**°**C for 5 min; 40 cycles at 95**°**C for 30 secs, 50**°**C for 1 min, 72**°**C for 1.5 min; and a final extension at 72**°**C for 10 min. Resulting PCR products were diluted 1:10 and used as template for second round PCR using conditions as per first-round, except for an annealing temperature of 55**°**C. PCR products were visualized using agarose gel electrophoresis and subsequently cloned into a pGEM-T easy cloning vector (Promega, Australia). Blue/white colonies were screened using standard techniques and a test digestion with endonuclease restriction enzyme *Eco*R1 was carried out to confirm insertion. Positive clones were sequenced in both directions using Sanger sequencing (AGRF).

### 2.6 Sequence alignment and phylogenetic analyses

A codon-matched nucleotide sequence alignment of 85 agnathan (jawless) and gnathostome (jawed vertebrate) opsin coding regions, ranging from lampreys to mammals, was generated by ClustalW (Higgins et al., 1996) and manually manipulated to refine the accuracy of cross-species comparison. Specifically, the alignment incorporated the opsin sequences of five visual photopigments expressed in the retina of *T. rugosa* (sleepy lizard) (Accession numbers: to be added upon acceptance) compared to those species listed in the phylogenetic tree. All five opsin classes were included, with several vertebrate ancient (VA) opsin sequences used collectively as an outgroup given that this opsin type is a sister clade to all five visual photopigment classes. Phylogenetic analyses of 1000 replicates were conducted in MEGA11 (Tamura et al., 2021), with evolutionary histories being inferred by using the Maximum Likelihood method and General Time Reversible model (Nei and Kumar, 2000). The percentage of trees in which the associated taxa clustered together is shown next to the branches. Initial trees for the heuristic search were obtained by applying Neighbour-Joining and BioNJ algorithms (Saitou and Nei, 1987) to a matrix of pairwise distances estimated using the Maximum Composite Likelihood (MCL) approach (Tamura and Nei, 1993). The tree was drawn to scale, with branch lengths measured in the number of substitutions per site. A total of 903 positions was present in the final dataset, with all positions with less than 95% site coverage being eliminated. That is, fewer than 5% alignment gaps, missing data, and ambiguous bases were allowed at any position.

### 2.7 Histological preparations

After completion of electrophysiological recordings, animals were euthanized with an intracelomic injection of sodium pentobarbital (Lethabarb, Virbac, Australia; dosage: 200 mg/kg). The dorsal region of the eye was cauterized prior to enucleation to allow for easy orientation. Once removed, the eye was opened with a small incision at the limbus using a scalpel blade and a small cut was made in the dorsal retina to maintain orientation. The cornea, lens and vitreous were removed and the eyecup preserved in 4% paraformaldehyde (PFA) in 0.1 M phosphate buffer (PB, pH 7.2-7.4) for both immunohistochemical analyses and for assessing retinal topography. Eyes were fixed in PFA for 24 h, then stored in 0.1 M PB plus 0.1% sodium azide. Radial cuts were made to relieve tension across the hemisphere of the eyecup and allow the retina to be flattened for wholemounts. The sclera and retinal pigment epithelium were removed to expose the retina.

### 2.8 Sampling of photoreceptor subtypes

Once the retinae were immunohistochemically labeled (see **Supplementary methods**), or simply mounted in glycerol, their outline was digitized using an X4.0 NA 0.17 objective lens, a motorized stage (MAC200; Ludl Electronics Products) and StereoInvestigator software (MBF Bioscience). The optical fractionator probe developed by West et al. (1991) and adapted by Coimbra et al. (2009) was used to sample the photoreceptor and retinal ganglion cell layer neurons.

A counting grid of a predetermined size (**Table S2**) was superimposed onto the retina with a randomized starting location. Each grid point that fell within the retinal outline represented a sampling location where a fraction of the area represented by that grid point was sampled using a sampling frame of predetermined size (**Table S2**). The number of cells marked within each sampling frame was extrapolated to estimate the number of cells located within the associated grid location. Total cell numbers in the retina were estimated by summing all grid locations (see **Error! Reference source not found**.).

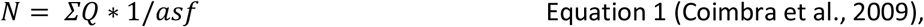

where *asf* is the area sampling fraction, ΣQ is the sum of markers counted within a frame and N is the total number of cells estimated within the area represented by the grid.

A total of six retinae was used to sample retinal neurons in *T. rugosa*. Three retinae were labeled with rho-4D2 (i.e., RH1 opsin) and three retinae were labeled with sc-14363 (i.e., SWS1 opsin). Retinae labeled with rho-4D2 were also used to investigate the overall distribution and numbers of photoreceptors by counting and mapping all photoreceptor types, including those that were unlabeled (i.e., single and double cones).

### 2.9 Generation of topographic maps

Topographic maps of photoreceptor distribution were constructed using a custom written Matlab (Mathworks, USA) function to determine how the distribution of photoreceptor subtypes differed across the retina. The sampling locations, retinal outline and cell locations were extracted from the xml file generated by the StereoInvestigator software (MBF Bioscience, USA). A thin plate spline was fitted (second order polynomial, lambda = 0) across sampling locations and used to interpolate the cell density at 20 µm intervals across the retina (Garza-Gisholt et al., 2014; Hemmi and Grünert, 1999). The spline reduced the effect of outlier fluctuations in the sampling, while providing a high-resolution estimate of cell density across the retina. Cell density across the retina was visualized using a combination of color maps and contour lines, which facilitated the identification of areas of retinal specialization.

## 3 Results

### 3.1 Transmission of ocular media and oil droplets

The lens and cornea were clear with half-peak transmittance (λT_0.5_) reached at 359 nm. The cornea was transparent to shorter wavelength light with a λT_0.5_ of 307 nm (Figure 1 Error! Reference source not found.). Lens and cornea combined absorbed 50% of the light at 391 nm (Figure 1) resulting in attenuated of UV light compared to higher wavelength.

**Figure 1.**
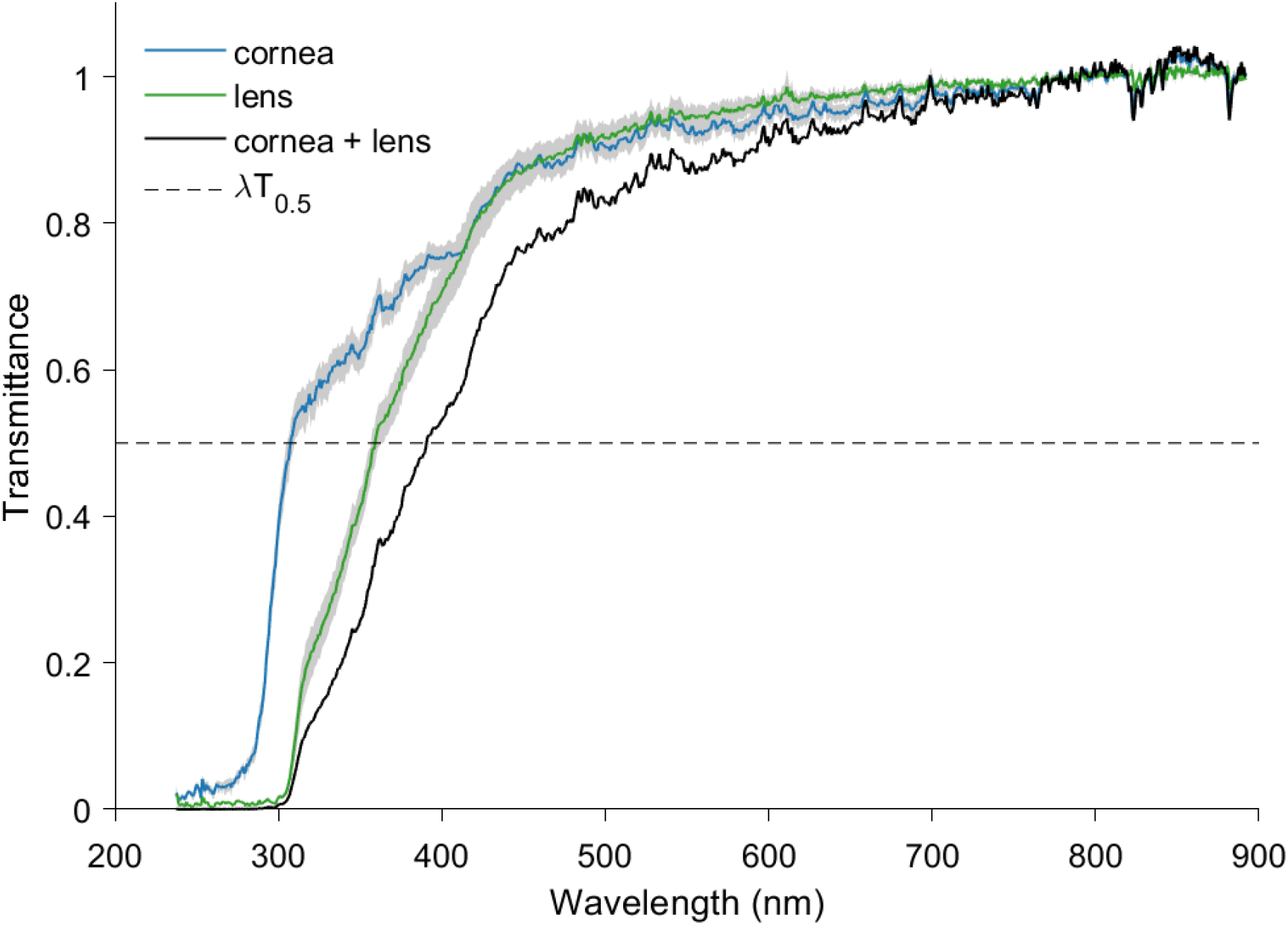
The lens is the major optical element which limits transmission at shorter wavelengths (λT_0.5_ at 359 nm). In contrast, the cornea transmits more light across the spectrum with λT_0.5_ at 307 nm. The combined optical elements achieve λT_0.5_ at 391 nm which would absorb more than half of all ultraviolet light entering the eye. The gray area around the transmittance curves of the cornea and lens indicates standard errors of the mean (s.e.m.). Individual transmittance curves were normalized to the average transmittance value from 840 nm to 690 nm. Cornea spectra were collected from six individuals while lens spectra were collected from four individuals.

Oil droplets appeared mainly of two types, transparent and pale-yellow with only a small number that could be reliably measured. The transparent oil droplet possessed equal absorptance throughout the spectrum, while the pale-yellow oil droplet had slightly increased absorptance below 430 nm (Figure 2). Absorptance profiles indicate that these two types of oil droplets are analogous to C2 (Figure 2 A) and C1 (Figure 2 B) oil droplets previously described in *Anolis valencienni* (Loew et al., 2002).

**Figure 2.**
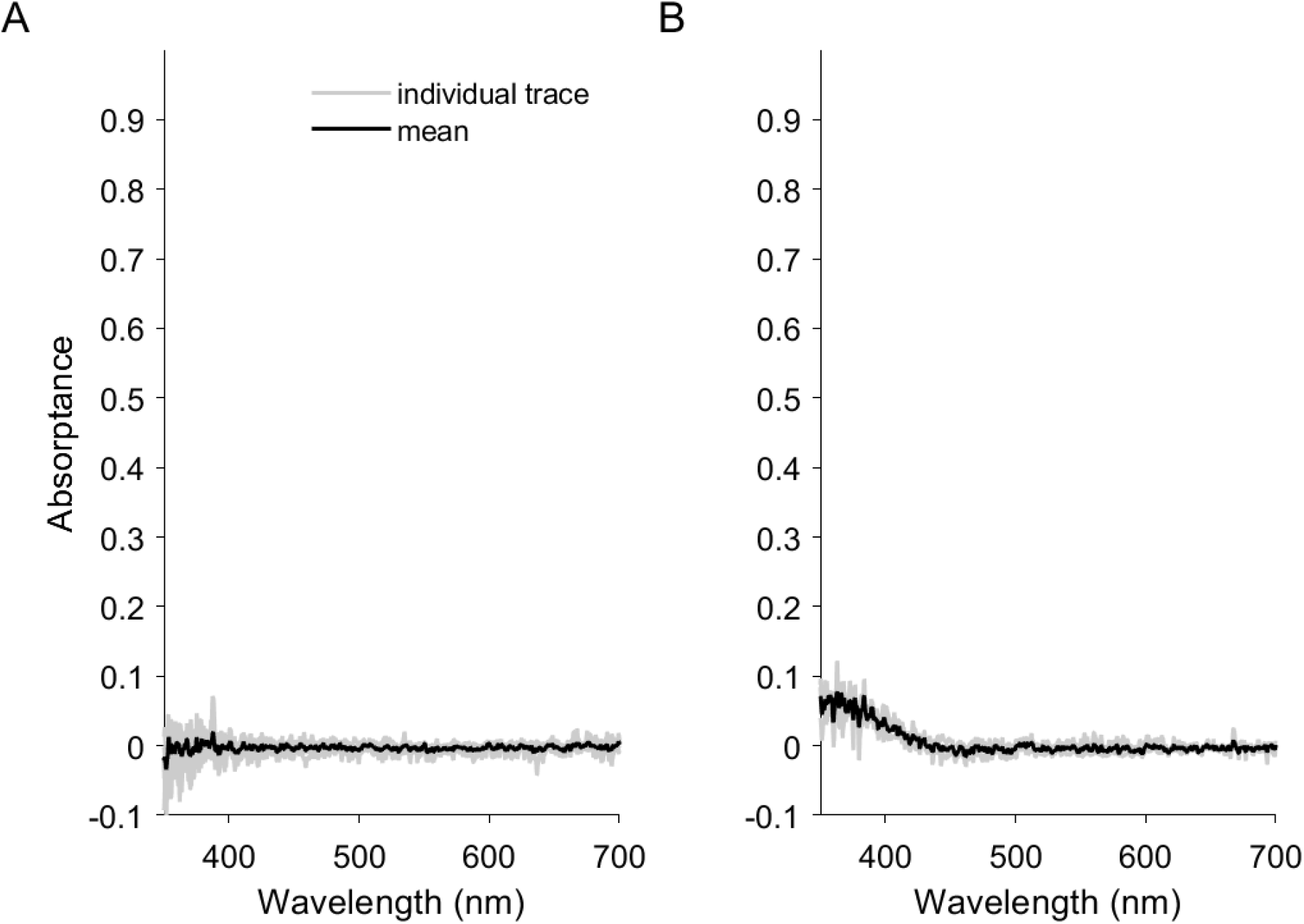
Absorptance spectra of *T. rugosa* oil droplets that are similar to the C2 (A) and C1 (B) oil droplets described in *Anolis valencienni* where they are associated with SWS1 and SWS2 photoreceptors, respectively (Loew et al., 2002). Mean absorptance (black line) is overlayed over absorptance traces of individual measurements (gray lines); n = 5 for A, n = 3 for B.

### 3.2 Spectral sensitivity

The spectral sensitivity of the sleepy lizard, *T. rugosa*, recorded with a stimulus frequency of 10 Hz peaked at 562±17 nm and was dominated by long-wavelength responses (Figure 3, blue line). The full width at half maximum (FWHM) of the spectral sensitivity curve was 135 nm ranging from 475 nm to 610 nm. This is much wider than the spectral sensitivity curves of eight other diurnal lizards (range 107 nm to 118 nm), even though they all peak in the same part of the spectrum (Figure 3). The spectral sensitivity of *T. rugosa* is broadened relative to other diurnal lizards due to increased sensitivity at shorter wavelengths. However, unlike in *P. broadleyi*, a secondary peak in the ultraviolet region was lacking.

**Figure 3.**
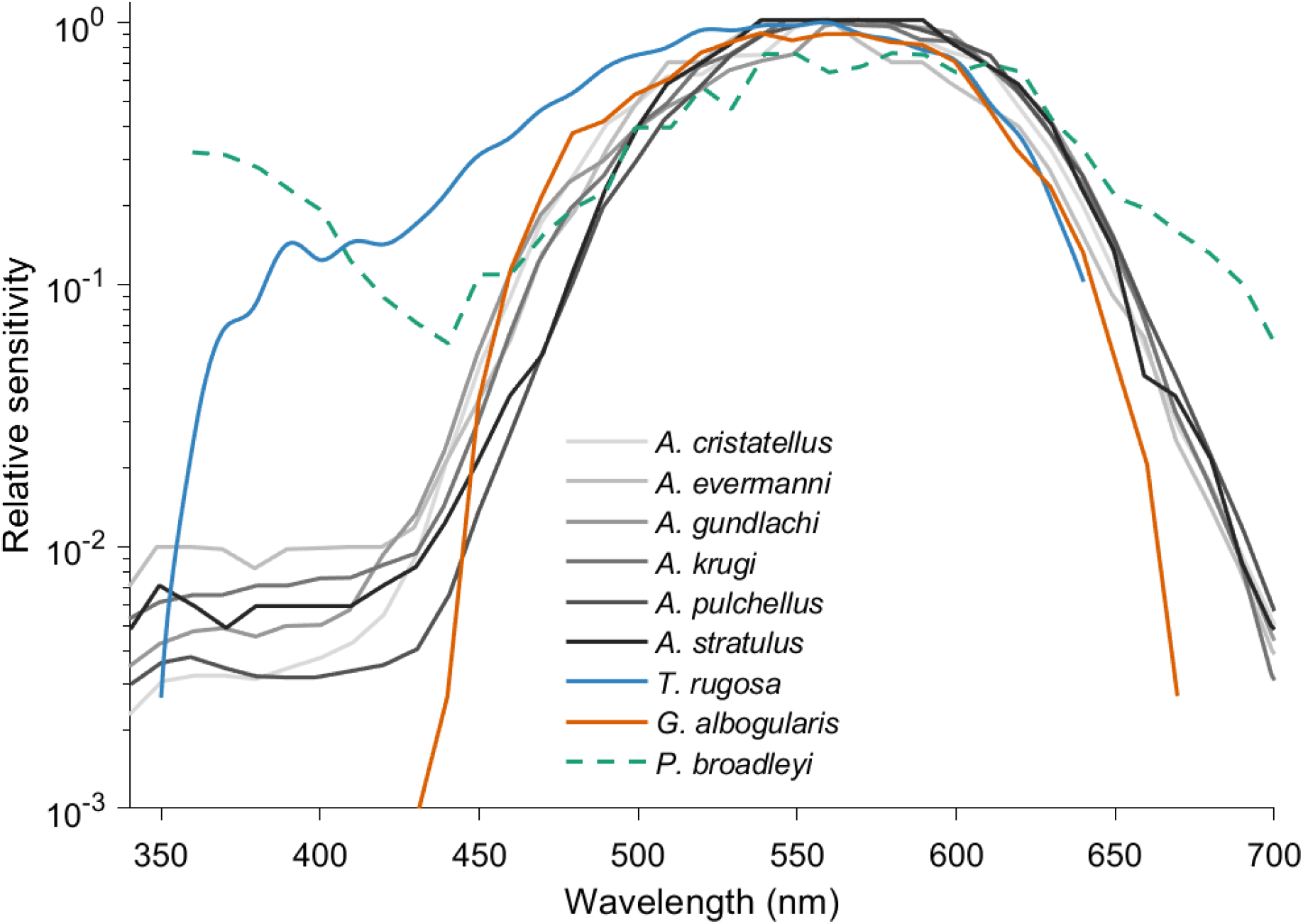
The spectral sensitivity of the sleepy lizard at a stimulus frequency of 10Hz peaks at 560 nm with a strong shoulder extending to about 390 nm (blue line). Sensitivity to short wavelength is much higher than in anole lizards and diurnal geckoes, where comparable stimuli were used (10-13Hz) (Fleishman et al., 1997, Ellingson et al., 1995). Although brief monochromatic flashes were used instead of flicker photometry to measure the spectral sensitivity of *P. broadleyi*, both it and *T. rugosa* possess much higher sensitivity to short wavelength light than other diurnal lizards. However, unlike in *P. broadleyi*, sensitivity to ultraviolet light drops dramatically in *T. rugosa*. Spectral sensitivity curves were plotted on a log scale here unlike in other plots to provide a clearer comparison to data gathered from Fleishman et al. (2011). Spectral sensitivity curves were normalized to the peak sensitivity.

Changing the flicker rate of the stimulus significantly changed the contributions of short-wavelength photoreceptors to the spectral sensitivity of the eye. The ratio of short-to long-wavelength sensitivity was significantly higher at 3 Hz than at 30 Hz (p<0.001, df = 19), indicating that photoreceptors sensitive to shorter wavelengths are contributing significantly more to the overall response at lower temporal frequencies. Compared to the standard 10Hz stimuli, a 3 Hz stimulus widened the spectral sensitivity curve (158 nm, Figure 4) by 23 nm while a 30 Hz stimulus narrowed the curve (129 nm, Figure 4) by 6 nm.

**Figure 4.**
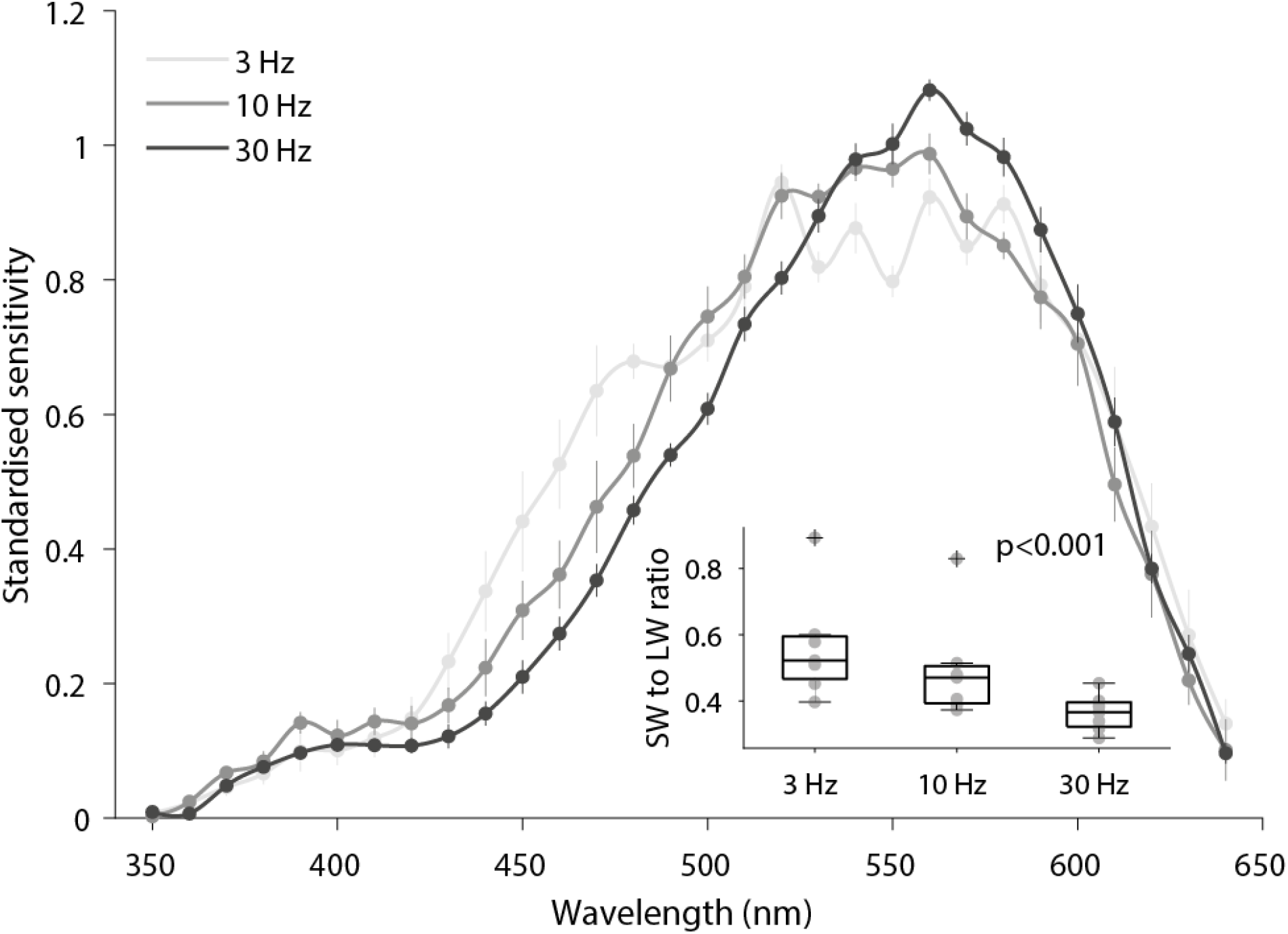
The spectral sensitivity (mean ± s.e.m.) of the eye of *T. rugosa* is narrower at higher temporal frequencies. The ratio of short-wavelength (<530 nm) to long-wavelength (>530 nm) sensitivity changed significantly across temporal frequencies with greater short wavelength photoreceptor contributions at lower temporal frequencies. A thin plate spline was fitted to ERG data with a lambda of 0. Spectral sensitivities were collected from 7 individuals for each condition. Boxplot limits in the inset denote 25^th^ and 75^th^ percentile and area between whiskers span ±2.7 standard deviations, datapoints beyond this area are considered outliers and illustrated with a plus sign. Individual spectral sensitivity curves were normalized to the average spectral sensitivity across conditions through a linear fit.

To reveal the spectral properties of the photoreceptor populations that could be contributing to this significant change, we used a monochromatic light at 550 nm. This selectively suppressed the contribution of LWS photoreceptors by reducing stimulus contrast around 550 nm and partially adapting photoreceptors that are very sensitive to the monochromatic light. At 10 Hz the addition of the 550 nm light shifted the peak sensitivity to 460 nm (p = 0.003, df = 7, Figure 5). Under the monochromatic light, varying the temporal frequency of our stimulus significantly changed the spectral sensitivity curve (p<0.001, df = 18, Figure 6). At slow flicker frequencies (3 Hz), the spectral sensitivity curve peaks at 470 nm and has a narrow bandwidth (72 nm). In contrast, at fast flicker frequencies (20-30Hz), the spectral sensitivity curve was dominated by photoreceptor responses with a peak sensitivity around 550 nm (Figure 6).

**Figure 5.**
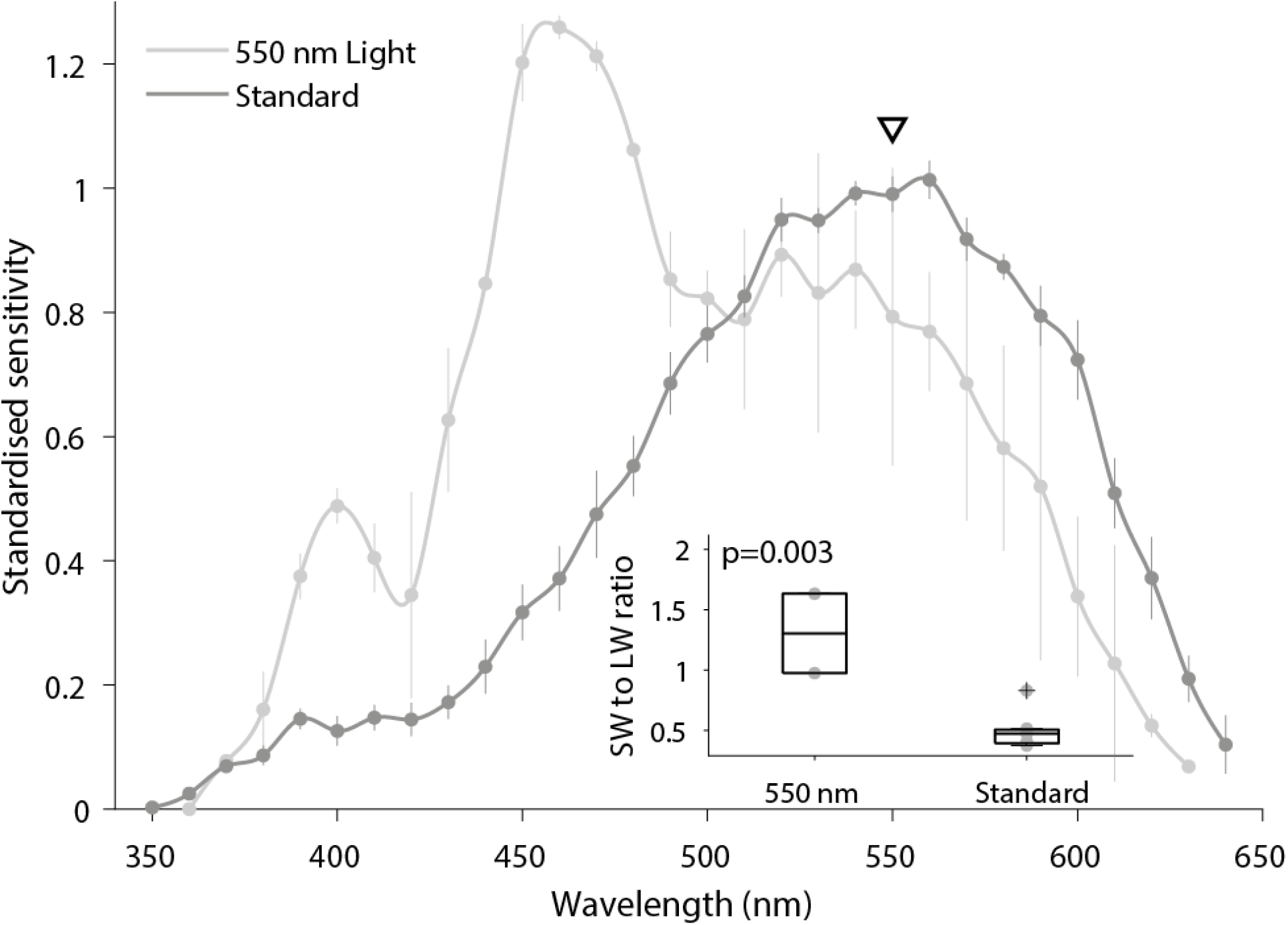
A monochromatic light at 550 nm used to suppress LWS responses, revealed contributions by a short-wavelength-sensitive cell population with a pronounced peak at 460 nm. The short-wavelength to long-wavelength sensitivity ratio is significantly higher with 550 nm light the standard 10 Hz stimulus. The black triangle indicates the spectral location of the 550 nm light. Sample size for conditions the standard and 550 nm light stimulus were seven and two, respectively. Boxplot limits in the inset denote 25^th^ and 75^th^ percentile and area between whiskers span ±2.7 standard deviations, datapoints beyond this area are considered outliers and illustrated with a plus sign. Individual spectral sensitivity curves were normalized to the average spectral sensitivity across conditions through a linear fit.

**Figure 6.**
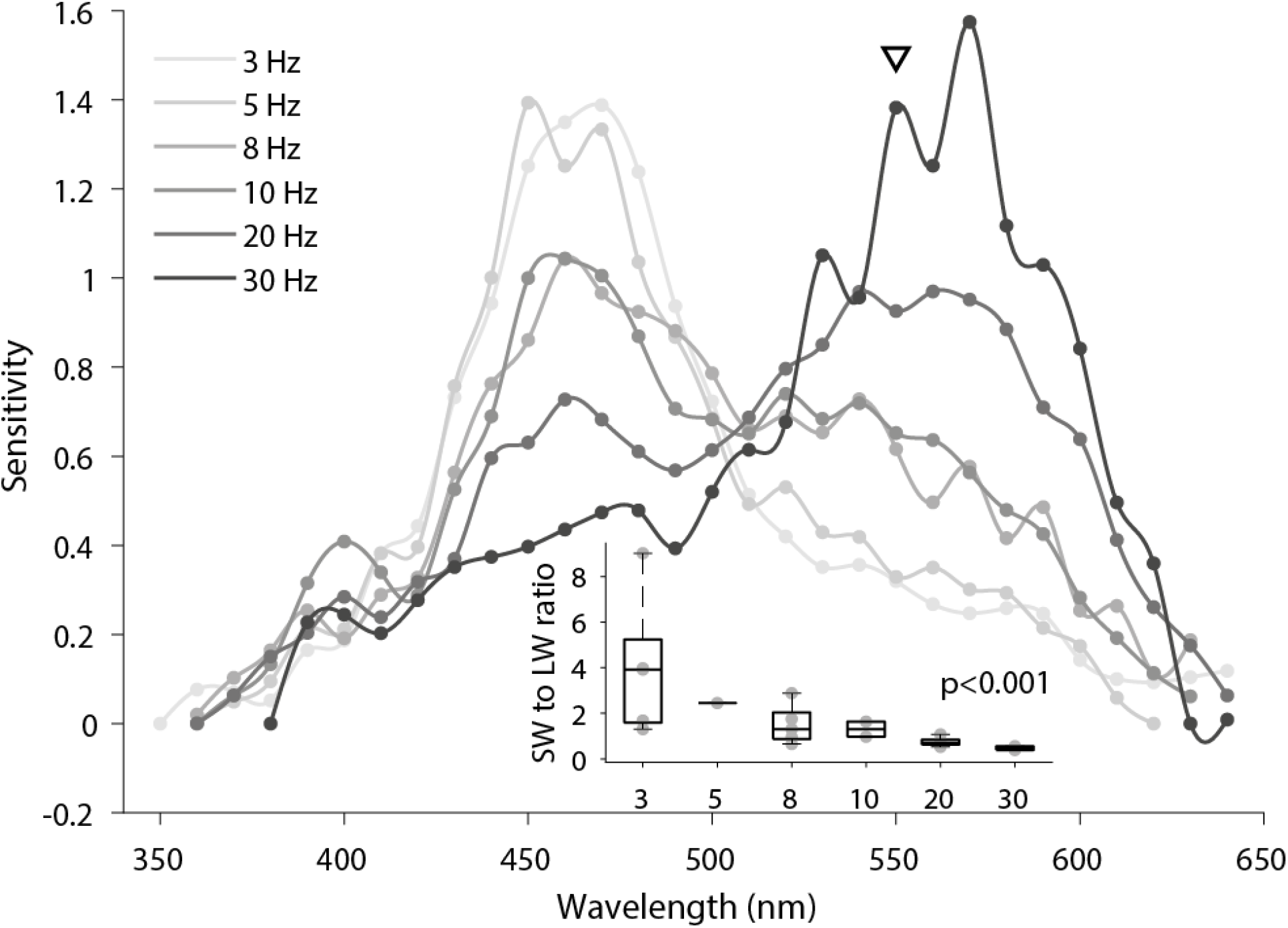
Applying a 550 nm monochromatic light while simultaneously lowering the temporal frequency of the stimulus, reveals a narrow spectral sensitivity curve likely dominated by a single photoreceptor subtype with peak sensitivity at 470 nm and tuned to low temporal frequencies. A thin plate spline was fitted to the ERG data with a lambda of 0. The black triangle indicates the spectral location of the 550 nm monochromatic light. Five individuals were sampled for 3Hz, 8Hz and 20 Hz, two individuals for 10Hz and 30Hz, and one individual for 5Hz flicker rates. Inset x-axis indicates the temporal frequency of experimental group, Hz unit was omitted for greater figure clarity. Boxplot limits in the inset denote 25^th^ and 75^th^ percentile and area between whiskers span ±2.7 standard deviations, datapoints beyond this area are considered outliers and illustrated with a plus sign. Individual spectral sensitivity curves were normalized to the average spectral sensitivity across conditions through a linear fit.

### 3.3 The reflectance of blue tongued lizards

The blue tongues of lizards reflect strongly at short wavelengths with a primary peak at approximately 320 nm and a secondary peak at approximately 460 nm (Figure 7, (Abramjan et al., 2015)). The pink tongue of *C. zebrata* has a similar bimodal reflectance profile but the balance of reflectance between short and long wavelength is shifted towards longer wavelength in *C. zebrata* (Figure 7).

**Figure 7.**
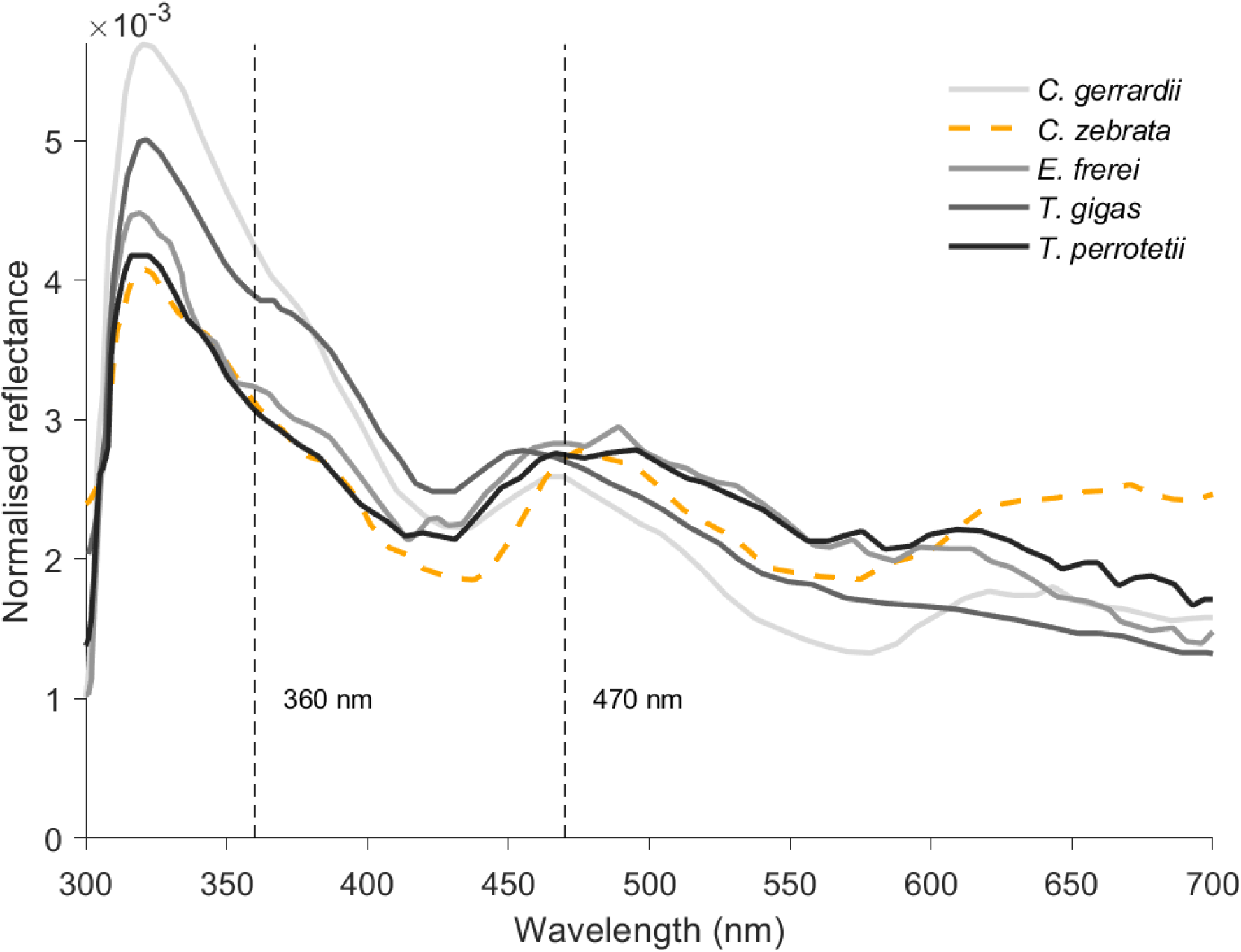
Normalised reflectance of pink (orange dashed line) and blue tongues (gray to black lines) of lizards adapted from Abramjan et al. (2015). Blue tongues have a higher normalised reflectance between 360 nm and 470 nm than the pink tongue of *Corucia zebrata*. Reflectance curves have been normalised to their respective integrals. Thin vertical black dashed lines indicate the precise location of 360 nm and 470 nm on the x-axis.

### 3.4 Opsin sequences

Five visual opsin genes were found to be expressed in the retina of the sleepy lizard. Sequence alignment and phylogenetic analyses confirmed them to be true orthologues of *LWS, SWS1, SWS2, RH2* and *RH1* opsin genes identified in other vertebrates (Figures 8 and Table S1). Upon closer inspection, it was possible to examine 30 out of the 34 known tuning sites across the five opsins genes and out of these only one tuning site (in the SWS2 opsin) differed from those in the green anole (*Anolis carolinensis*). There is a high sequence similarity between the opsins of the sleepy lizard and the green anole (Figure S1).

**Figure 8.**
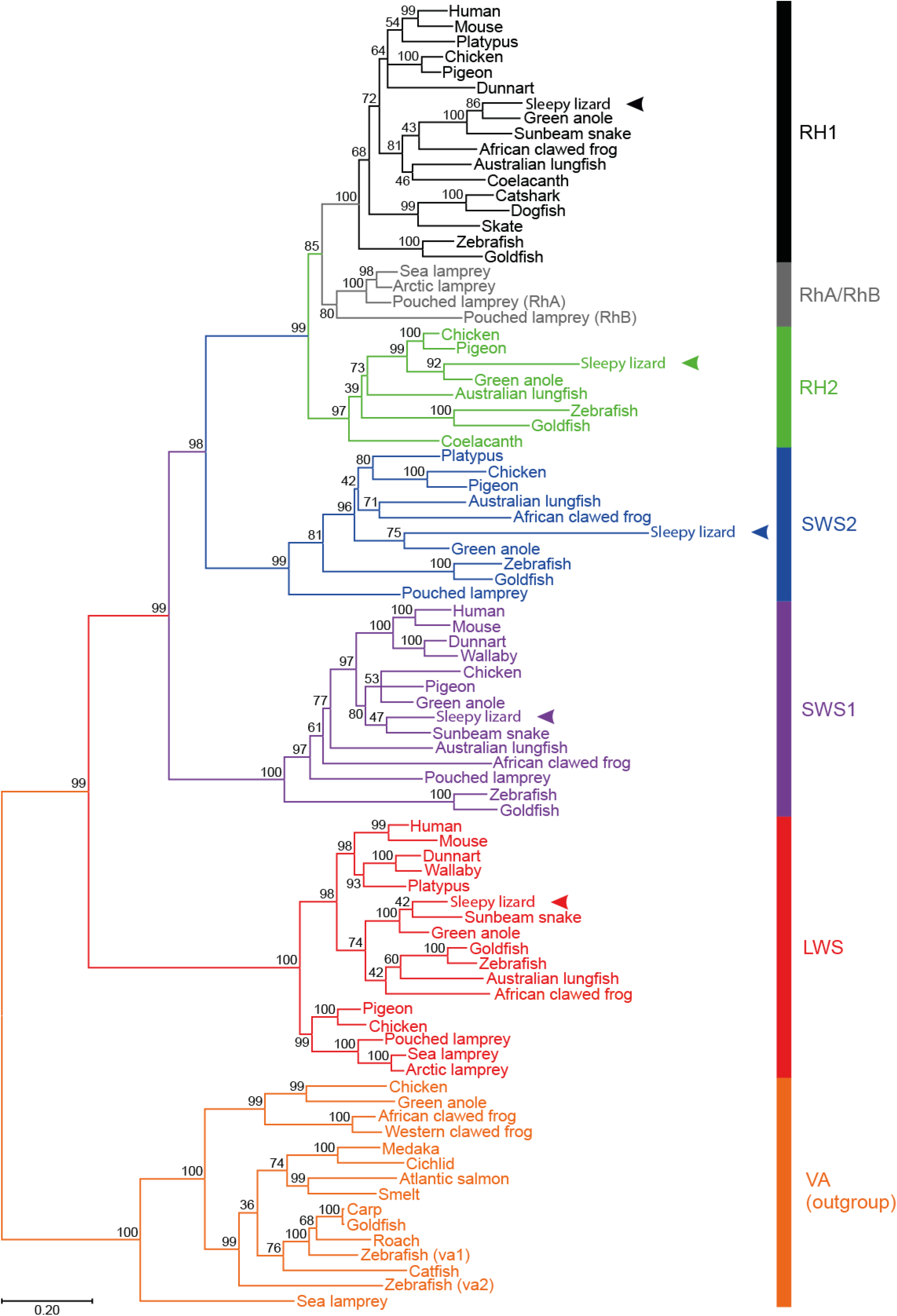
The evolutionary history was inferred by using the Maximum Likelihood method and General Time Reversible model (Nei and Kumar, 2000). The tree with the highest log likelihood (−38890.45) is shown. The percentage of trees for which the associated taxa clustered together is shown next to the branches. Initial tree(s) for the heuristic search were obtained automatically by applying Neighbour-Join and BioNJ algorithms (Saitou and Nei, 1987) to a matrix of pairwise distances estimated using the Maximum Composite Likelihood (MCL) approach (Tamura and Nei, 1993), and then selecting the topology with superior log likelihood value. A discrete Gamma distribution was used to model evolutionary rate differences among sites (5 categories (+*G*, parameter = 0.9476)). The rate variation model allowed for some sites to be evolutionarily invariable ([+*I*], 10.93% sites). The tree is drawn to scale, with branch lengths measured in the number of nucleotide substitutions per site (indicated by the scale bar). This analysis involved 85 nucleotide sequences, where the five *T. rugosa* visual photopigment genes (*LWS, SWS1, SWS2, RH2* and *RH1*; indicated by arrowheads) have the following Accession Numbers: TBD upon acceptance. Codon positions included were 1st+2nd+3rd+Noncoding. All positions with less than 95% site coverage were eliminated, i.e., fewer than 5% alignment gaps, missing data, and ambiguous bases were allowed at any position (partial deletion option). There was a total of 903 positions in the final dataset. Evolutionary analyses were conducted in MEGA11 (Tamura *et al*., 2021).

### 3.5 Retinal topography

Five photoreceptor counts were made and mapped separately, including the total photoreceptors, single cones, double cones, RH1 cones and SWS1 cones (the latter two were isolated immunohistochemically, Figure 9 A-B). The total photoreceptor population comprised 77.6% single cones and 22.4% double cones (Table 1). No labelling with antibodies specific to RH1 or SWS1 was observed in double cones, suggesting that principal and accessory members of the double cones contain either SWS2, RH2 or LWS opsins. RH1 photoreceptors comprised 22.8% of single cones and 17.7% of all photoreceptors. In contrast, the number of SWS1 cones was significantly lower at 8.5% of single cones and 6.6% of the total photoreceptor population (Table 1).

**Table 1.**
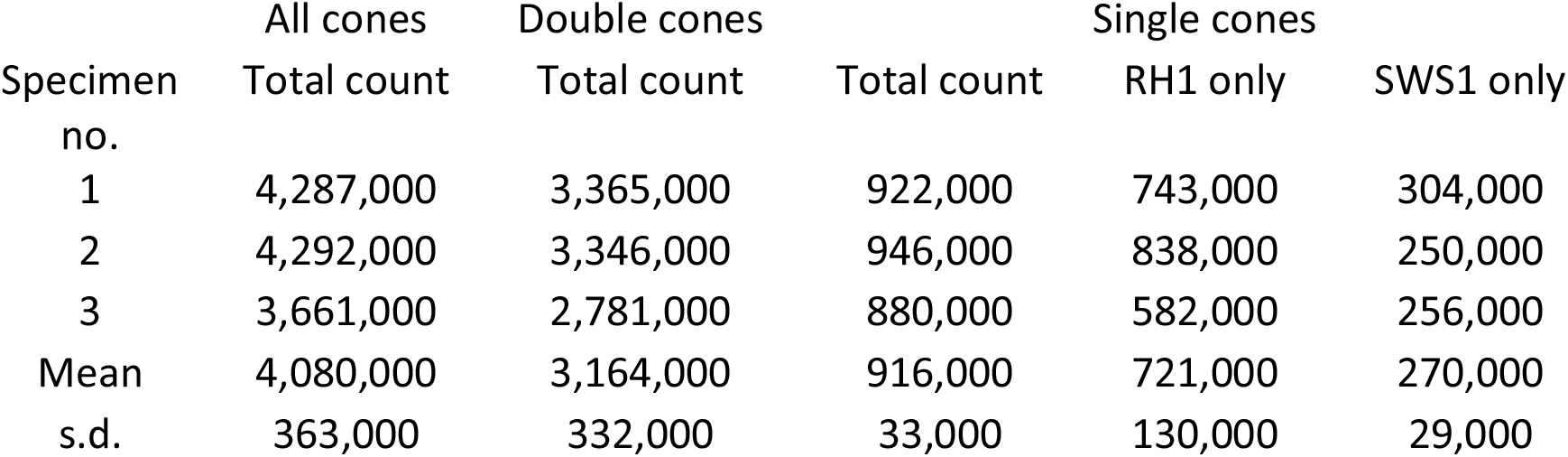
Photoreceptor counts in the retina of *T. rugosa* showed that single cones made up 77.6% of all photoreceptors, with the rest being double cones. Of these single cones, 22.8% were identified as RH1 cones, with the presence of 8.6% SWS1 cones, although the latter made up 6.6% of the total photoreceptor population.

**Figure 9.**
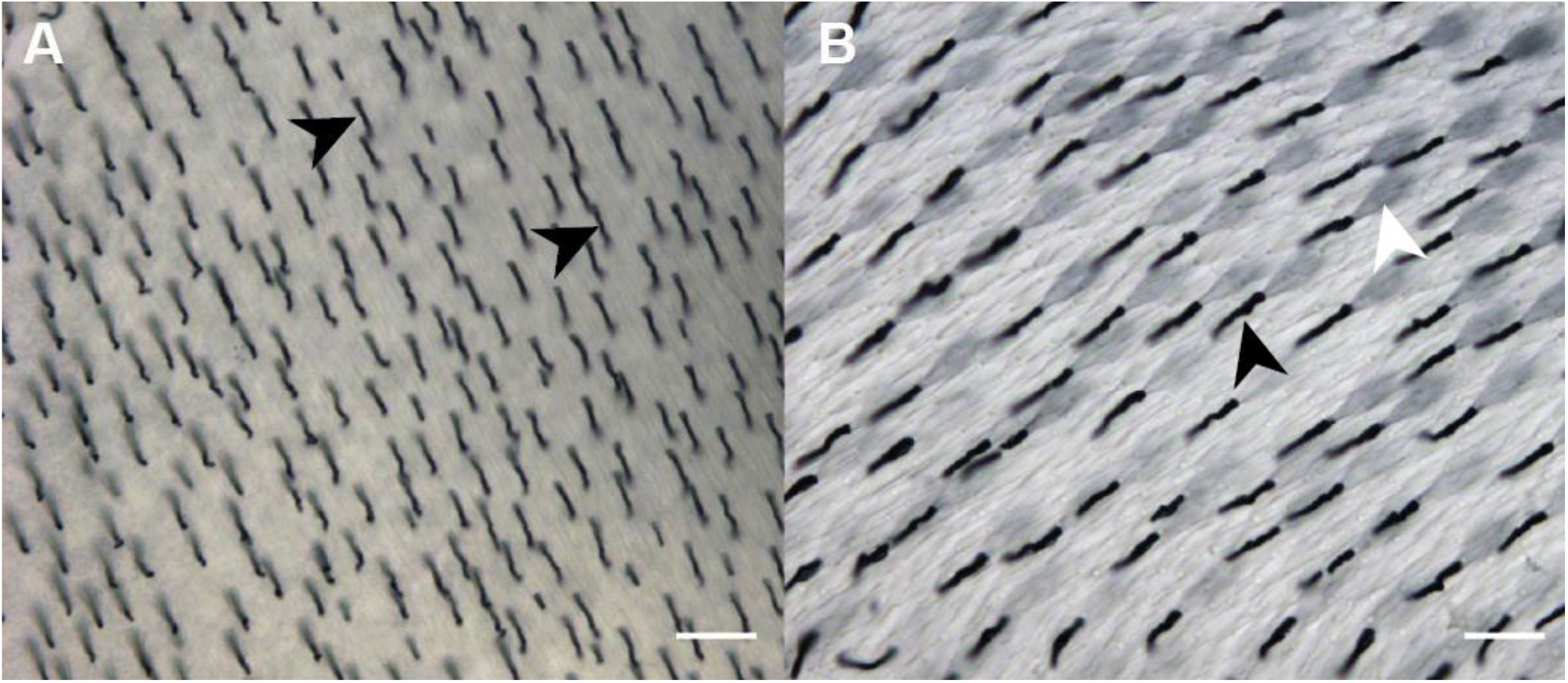
Rho-4D2 and sc-14363 stained outer segments of RH1 (A) and SWS1 (B) in the retina of Tiliqua rugosa. Black arrowheads indicate the outer segments, and the white arrowhead indicates the inner segment of photoreceptors. Scale bars = 20 µm.

All single cones and double cones had similar distributions across the retina. The density increased steadily towards the retinal center forming an area centralis. There was a slight asymmetry with a higher number of photoreceptors in the ventral compared to the dorsal retina (Figure 10 A-D). Although no clear horizontal visual streak was observed, relatively high densities (between 35,000 to 55,000 photoreceptors/mm^2^) were maintained across the naso-temporal axis of the retina at the retinal meridian (Figure 10 A-D). SWS1 single cones also adopted a concentric increase in density towards the central retina but had a much shallower gradient with densities ranging from 2,000 to 5,000 SWS1 photoreceptors/mm^2^ in the retinal center (Figure 10 C).

**Figure 10.**
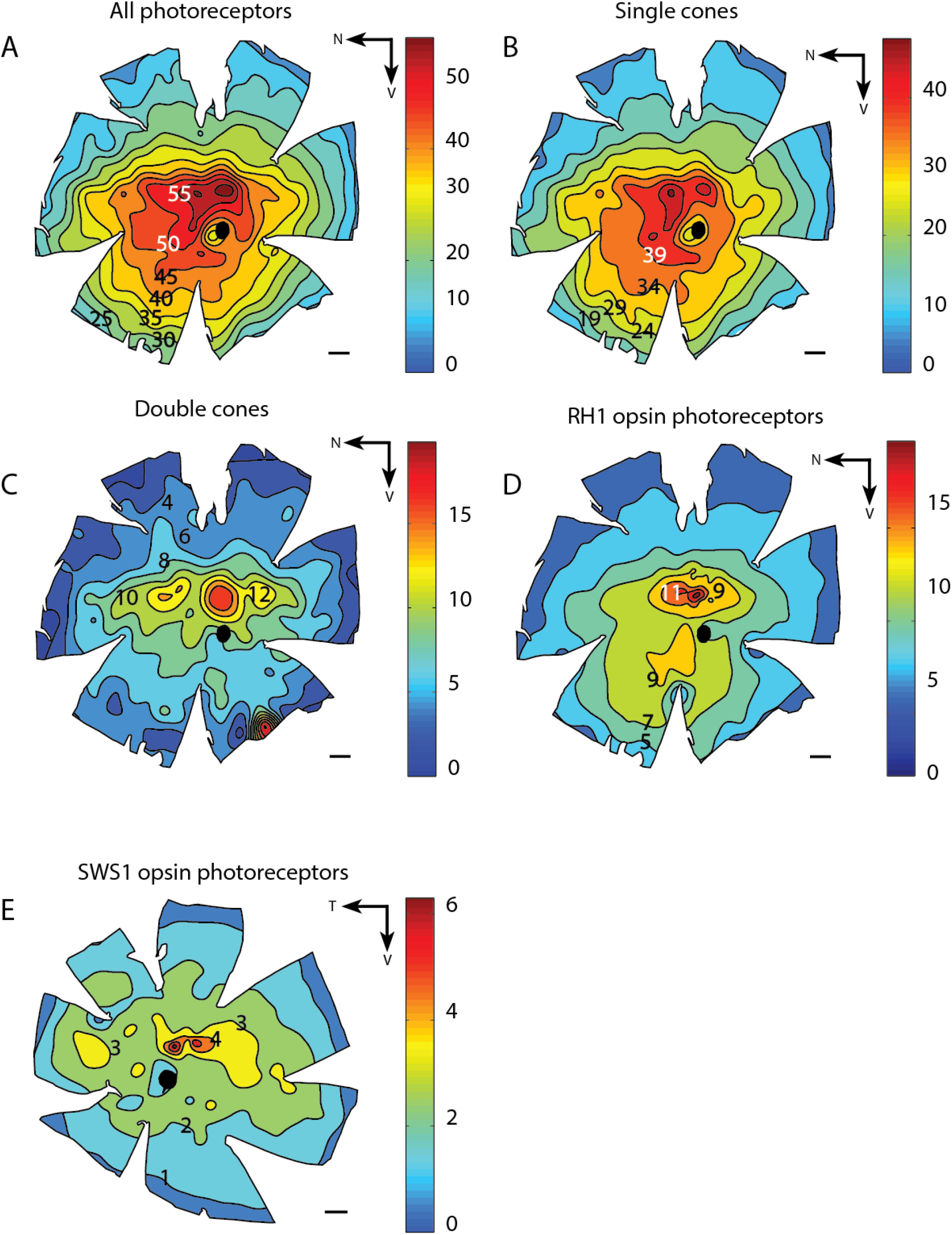
All photoreceptor populations showed peak densities in the retinal center, with an area centralis and ventral wedge of high photoreceptor density (A), single cones (B), double cones (C) and RH1-expressing cones (D). SWS1-labeled photoreceptors (E) were fewer in number and showed a shallower slope of density increase towards the center. N, nasal retina; T, temporal retina; V, ventral retina. Scale bars indicate 1 mm across all maps. Black circles in the middle of the maps indicate the optic nerve head. Iso-density lines and colored bars indicate density in cells/mm2 (X1000).

## 4 Discussion

Our findings reveal that *T. rugosa* has an elevated sensitivity at shorter wavelength which is distinct from other diurnal lizards. Here, we discuss potential drivers for these differences such as spectral tuning and photoreceptor abundance, their ecological benefits, and future research directions.

### 4.1 Sensitivity to ultraviolet light differs from other diurnal lizards

A close look at the ultraviolet portion of the spectral sensitivity curves of diurnal lizards studied to date (Figure 3) reveals that a secondary peak occurs between 350 nm and 360 nm followed by a slight reduction in sensitivity at 370 nm to 380 nm. This is typically associated with the peak sensitivity of SWS1 photoreceptors which, in diurnal lizards, typically ranges from 358 nm to 367 nm. Influences from the beta peak of LWS and MWS photoreceptors are likely minimal or non-existent in the short wavelength region due to their association with oil droplets which typically absorb most/all light below ∼470 nm as described in the anole lizard retina (Loew et al., 2002). In contrast to most diurnal lizards, the cordylid lizard, *Platysaurus broadleyi*, shows elevated sensitivity to the ultraviolet at 360 nm which has been associated with an increased abundance of UVS photoreceptors (Figure 3, green dashed line, Fleishman et al. 2011). However, in the *T. rugosa*, there appears to be no UV peak, but a sharp drop after 380-390nm. There is no conspicuous peak in sensitivity to ultraviolet light even when using stimuli that should simultaneously suppress LWS contributions while increasing the contributions of SWS photoreceptors (550 nm monochromatic light + low temporal frequencies, Figure 6). This suggests that despite the presence of SWS1 which is typically associated with UVS photoreceptors in diurnal lizards, UVS photoreceptors may be absent in *T. rugosa*. This could be caused by a shift of the spectral sensitivity of SWS1 photoreceptors towards longer wavelengths.

### 4.2 Red-shifting the spectral sensitivity of the SWS1 photoreceptor

#### 4.2.1 Single site amino acid substitution?

Large shifts in the spectral sensitivity of SWS1 photoreceptors from ultraviolet to violet are commonly associated with single site amino acid substitutions at site 86 of the SWS1 photopigment (Hauser et al., 2014). In freshwater (*Helicops modestus*) and sea (*Aipysurus spp*., *Epiocephalus spp*., *Hydrophis spp*.) snakes, the amino acid phenylalanine is substituted for valine, serine, cysteine or tyrosine at site 86 of the SWS1 photopigment, thereby shifting its spectral sensitivity towards longer wavelengths (Hauzman et al., 2017; Simões et al., 2020). All diurnal lizards studied to date, possess phenylalanine at site 86, which results in UV-sensitive photopigments, however, in contrast to aquatic snakes, no amino acid substitutions have yet been reported. While we have been unable to confirm the amino acid identity at site 86 of the SWS1 photopigment in *T. rugosa*, the conspicuous lack of UV sensitivity in the ERGs suggest that phenylalanine has been substituted for another amino acid to produce violet-sensitive (VS) photoreceptors instead of UVS photoreceptors.

#### 4.2.2 Alternative spectral tuning mechanisms and why they are less likely

Mechanisms such as changes from an A_1_ photopigment to an A_2_ photopigment, and the filtering effects of ocular media and oil droplets, may also shift the spectral sensitivity of retinal photoreceptors. However, we describe below, how their potential to shift the spectral sensitivity of a hypothetic UVS photoreceptor are limited and would not be congruent with the lack of a peak in sensitivity in the ultraviolet region of Figure 6 for low temporal frequency stimuli.

Transitions from A_1_ to A_2_ chromophores are commonly used in fish to shift the spectral sensitivity of their photopigments to longer wavelengths (Bowmaker, 1995; Whitmore and Bowmaker, 1989), however, in diurnal lizards, this is relatively rare with the adoption of pure A_2_ photopigments only observed in the green anole, *A. carolinensis* (Kawamura and Yokoyama, 1998). SWS, MWS and LWS photopigments of the green anole were shifted towards longer wavelengths, but the UVS photoreceptor spectral sensitivity remained unchanged by the adoption of the A_2_ chromophore (Loew et al., 2002). In the unlikely event that *T. rugosa* uses pure A_2_ chromophores, we therefore expect little to no effect on the spectral sensitivity of potential UVS photoreceptors due to high sequence similarity of photopigments between green anoles and *T. rugosa*.

Oil droplets associated with SWS1 photoreceptors tend to be transparent in all measurements in lizards to date, and we have identified a transparent oil droplet likely to be similarly associated in *T. rugosa*. We therefore expect little to no effect of oil droplets on the SWS1 photoreceptors. In contrast to the oil droplets, the λT_0.5_ of the lens is long wave-shifted in *T. rugosa* (356 nm) compared to diurnal lizards (∼320 nm, (Pérez i de Lanuza and Font, 2014)) and would absorb more than half of the UV light entering the eye. On their own, the cornea and lens of *T. rugosa* combined would shift the λ_max_ of a typical diurnal lizard SWS1 photoreceptor from ∼365 nm to 377 nm, however, this would have the effect of substantially reducing the photon capture ability of such a SWS1 cone (see Figure 6).

### 4.3 Short wavelength sensitivity is greatly enhanced by multiple factors

#### 4.3.1 The role of SWS2 photoreceptors

Despite several lines of evidence suggesting that *T. rugosa* do not possess a UVS photoreceptor, they are up to an order of magnitude more sensitive than other diurnal lizards between 360 and 530 nm (Figure 3) due to the broadness of its spectral sensitivity curve. The effect of adjusting the temporal frequency of our stimulus on the width of the spectral sensitivity curve suggests that the wider spectral sensitivity curve of *T. rugosa* is partially mediated by a population of photoreceptors that favor low temporal frequencies (Figure 4). Peak sensitivities between 450-470 nm under 550 nm monochromatic light at low temporal frequencies suggest that these are SWS2 photoreceptors. This is supported by the λ_max_ of SWS2 photoreceptors at similar wavelengths in other lizards (Osorio, 2019) and the well-established high sensitivity of this photoreceptor type to low frequency stimuli across taxa including tiger salamanders, goldfish, ants, and primates (Howlett et al., 2017; Ogawa et al., 2015; Rieke and Baylor, 2000; Skorupski and Chittka, 2010; Tailby et al., 2008). An increased abundance of SWS2 photoreceptors in *T. rugosa* could explain the enhanced short-wavelength sensitivity observed here. This is different to the suspected increased abundance of SWS1 photoreceptors in *P. broadleyi* which is thought to enhance sensitivity to ultraviolet light but leaves a dip in sensitivity to blue light (∼450 nm) (Fleishman et al., 2011; Martin et al., 2015). However, it is clear that the SWS2 photoreceptors alone cannot explain the relatively broad spectral sensitivity curve of *T. rugosa* as the curve remains broader than that of other diurnal lizards even under conditions (30 Hz, Figure 4, Figure 6) where the response of SWS2 photoreceptors was greatly attenuated. It is worth noting that a red-shifted SWS1 opsin would have a greater overlap with the neighboring SWS2 opsin and would therefore increase the quantal catch of the eye where the spectra of the photoreceptors overlap.

#### 4.3.2 Oil droplets associated with MWS and LWS photoreceptors

Typically, MWS and LWS photoreceptors are associated with green (G), orange (O) or yellow (Y) oil droplets which absorb most of the light below 470 nm thereby reducing their contribution to short-wavelength sensitivity (Barbour et al., 2002; Crescitelli, 1972; Loew et al., 2002; Martin et al., 2015). However, our findings and the previous reports by New et al. (2012) suggest that *T. rugosa* only has two types of oil droplets, both of which transmit most of the light below 470 nm. New et al. (2012) further reported that pale-yellow oil droplets are associated with single cones and the principal member of the double cone, which are typically LWS photoreceptors (Crescitelli, 1972). This suggests that in *T. rugosa*, at least the LWS photoreceptor, if not both the MWS and the LWS photoreceptors, are associated with pale-yellow oil droplets. This would result in significantly greater sensitivity to shorter wavelength light and could partially explain the broad spectral sensitivity curve of *T. rugosa* relative to other diurnal lizards.

### 4.4 Insights from the abundance and distribution of photoreceptors

#### 4.4.1 Photoreceptor subtype abundance

The typical abundance of SWS1 photoreceptors in birds and mammals sits around 5-10% (Hart, 2001; Hunt and Peichl, 2014; Kram et al., 2010). Similar estimates have been observed in turtles (6%, *Pseudemys scripta*) and anole lizards (5%, *A. sagrei*) where the SWS1 opsin is maximally sensitive to UV (Fleishman et al., 2011; Grötzner et al., 2020). In *Platysaurus broadleyi* which possesses enhanced UV sensitivity, ERG data and transparent oil droplet counts suggests that 16.5% of photoreceptors are UVS photoreceptors, which are likely to contain the SWS1 opsin. In contrast, only 6.6% of photoreceptors in the retina of *T. rugosa* express SWS1 indicating that an increased abundance of SWS1 is not driving enhanced sensitivity to short wavelengths in this species. Instead, the abundance of SWS1 photoreceptors is similar to the broadly conserved percentages of SWS1 photoreceptors found across taxa.

SWS2 photoreceptors, identified through oil droplet counts in birds and turtles, comprise 10-15% of the total population of photoreceptors, which is 1-1.5 times the number of SWS1 photoreceptors (Grötzner et al., 2020; Hart, 2001). Assuming that those proportions are preserved in *T. rugosa*, we can assume that at least 6.6-9.9% of photoreceptors will express SWS2 opsins or a higher percentage if the abundance of SWS2 photoreceptors are elevated in this species.

The presence of RH1 in cone like cells has been widely reported in other diurnal lizards and snakes, but this is the first time the proportion of RH1 cones have been accurately quantified over the entire retina of a squamate. Our finding that 17.7% of photoreceptors express the RH1 opsin, suggest that these transmuted cells (Walls, 1942) make a significant contribution to the response of the whole eye. While we successfully labelled and quantified the abundance of RH1 photoreceptors, it remains unknown what percentage of the photoreceptor population is comprised of RH2 photoreceptors.

While LWS photoreceptors were not labeled in this study, previous reports by New et al (2012) indicate that the LWS opsin is not co-expressed with the SWS1 or RH1 opsins in the retina of *T. rugosa*. The expression of the LWS opsin in the accessory and principal members of the double cones, as well as in a subset of single cones, is well established in diurnal lizards (Loew et al., 2002). This expression pattern is also present in birds and turtles suggesting a well-preserved feature likely repeated in the retina of *T. rugosa*. Given that double cones comprise 22% of the total photoreceptor population, it is highly likely that LWS photoreceptors are the most abundant photoreceptor subtype in the retina of *T. rugosa*. This would align well with the spectral sensitivity of the whole eye peaking at ∼560 nm observed in this study and in other studies of diurnal lizards (Ellingson et al., 1995; Fleishman et al., 1997).

#### 4.4.2 Photoreceptor subtype topographic distribution

The high densities of single cones and double cones which persist towards the ventral periphery of the retina suggest that the dorsal visual field is spatially sampled at a higher resolution than the ventral visual field of *T. rugosa*. This is likely, as the head of *T. rugosa* is very close to the ground with most ecologically relevant interactions likely to occur at eye level or within the dorsal visual field. Similar patterns which mirror our observations are found in the retina of artiodactyls, where height is coupled to the density of photoreceptors in the dorsal retina, with taller artiodactyls possessing higher densities of photoreceptors in their dorsal retina (Schiviz et al., 2008). However, unlike other types of photoreceptors mapped here, the SWS1 photoreceptors are rather uniformly distributed. Due to the low abundance of SWS1 photoreceptors, they may be less relevant to high acuity vision and therefore, they may not be under the same selective pressure which produces dorso-ventral asymmetries in the distribution of other photoreceptor types.

Up to now, the distribution of RH1 photoreceptors in squamates has remained unmapped. The dorsoventral asymmetry in the distribution of RH1 photoreceptors is not dissimilar to previous findings in passerine birds and various mammals (Coimbra et al., 2015; Kryger et al., 1998; Müller and Peichl, 1989). However, unlike in the passerines and raptors (Coimbra et al., 2015; Mitkus et al., 2017), the peak densities of RH1 photoreceptors coincide with the peak of all photoreceptors and there are no retinal areas where the RH1 opsin is not expressed. In *A. carolinensis*, RH1 photoreceptors are expressed even in the fovea whereas in bird foveas, RH1 photoreceptors are completely absent (Coimbra et al., 2015; McDevitt et al., 1993; Mitkus et al., 2017). While *T. rugosa* does not possess a fovea, the difference in the retinal distribution of these cells between birds and squamates suggest that RH1 photoreceptors play a different role in their respective visual systems (Schott et al., 2016; Zhang et al., 2006).

### 4.5 Ecological relevance of higher sensitivity to short wavelengths

*T. rugosa* belongs to a group of lizards that possess blue tongues, which they are suspected of using to deter predators and to avoid aggressive male-to-male interactions (Abramjan et al., 2015; Badiane et al., 2018). Despite the similar reflectance profile of blue and pink tongues, blue tongues tend to reflect relatively more short wavelength light (Figure 7) as the pigment in their tongues likely absorb longer wavelength light (Abramjan et al., 2015). Previous visual models report that blue tongues are more conspicuous to species of the *Tiliqua* genus than to their potential aerial predators (Abramjan et al., 2015), however, this assumes that *Tiliqua spp*. possess similar enhanced UV sensitivity to *P. broadleyi*. Instead, our findings show that *T. rugosa* possesses enhanced sensitivity over a broader region of the spectrum and may be missing UVS photoreceptors. While this may still make the blue tongue, which is common in this genus, more conspicuous to its conspecifics than its predators, it suggests that signal detection is fundamentally different to what has been previously reported.

## 4.6 Conclusion

The eye of *T. rugosa* is up to an order of magnitude more sensitive to short wavelengths than other diurnal lizards suggesting that the detection of short wavelenght light plays relatively important role in the ecology of this species. This enhanced sensitivity appears to be achieved by multiple factors which involve blue photoreceptors tuned to slow temporal frequencies, the absence of yellow and green oil droplets and potentially red-shifted SWS1 photoreceptors. While our findings demonstrate the effect of these factors on the overall sensitivity of the eye, the precise nature of how these mechanisms are being implemented in the retina of *T. rugosa* remain unclear. Additional studies should fully sequence the tuning sites of the SWS1 opsin, rigorously characterise the association between oil droplets and photorecepor types and accurately estimate the abundance of SWS2 photoreceptors While this enhanced sensitivity to short wavelengths may facilitate the detection of the blue tongue, which is commonly used as a conspecific signal, the relationship between the two remains unclear as other factors may also be responsible for the increased sensitivity to short wavelength observed in this species. Taken together with isolated studies in *P. broadleyi* and *Z. vivipara*, our findings suggest that the co-evolution of the lizard eye and the perception of a diverseity ofvisual signals may be more complex than previously proposed. As opposed to dermal UV-reflective patches, the blue tongue found in this species is a rare signalling feature constrained to species of the *Tiliqua* genus and a few other species of skinks. This offers a unique opportunity to dissect the phylogenetic and environmental pressures which shape the spectral sensitivity of diurnal lizards in understudied clades of lizards.

## 4.8 Supplementary Methods

### 4.8.1 Immunohistochemistry and antibody specificity

The use of immunohistochemistry to label photoreceptor subtypes in the retina of T. rugosa was limited to antibodies where high opsin specificity had been previously demonstrated. As such, only SWS1 and RH1 opsins could be successfully immunohistochemically labeled.

The sc-14363 goat polyclonal antibody (Santa Cruz Biotechnology, USA) was raised against the epitope of the human S cone opsin and has been successfully used to label SWS1 opsins in birds (Nießner et al., 2011). This is mainly due to the high similarity of amino acid sequences between the S opsin epitope of mammals and the SWS1 opsin epitope of birds (Hart and Hunt, 2007) and low epitope sequence identity to other opsins (Nießner et al., 2011). Hence, labeling of SWS1 cones by sc-14363 is highly specific and, in birds at least, shows no overlap with labeling by JH492 antibody that detects LWS-derived photopigments (Nießner et al., 2011).

The rho-4D2 mouse monoclonal antibody was raised against the N-terminus of bovine rhodopsin and has been successfully used to specifically label RH1 opsins in several lizard species, including the American chameleon (Anolis carolinensis), the chameleon Chameleo chameleon and the sleepy lizard, Tiliqua rugosa (McDevitt et al., 1993, New et al., 2012).

Retinae were incubated overnight without agitation to minimize tissue damage in a solution of 0.3% Triton X100 in 0.1 M PB (pH 7.2-7.4), 5% normal rabbit serum (rho-4D2) or 5% normal donkey serum (sc-14363) and the primary antibody (1:500 for both antibodies). Prior to being incubated with a biotinylated secondary antibody for 2 h, retinae were rinsed in 0.1 M PB buffer three times for 5 min each, before being transferred to a solution containing an avidin-biotin complex (Vectastain, ABC kit, Vector Laboratories, USA), incubated for 1 h, then rinsed three times (5 min each) in 0.1 M PB. Opsin immunoreactivity was visualized via a horseradish peroxidase (HRP) and H2O2 reaction as recommended in the peroxidase substrate kit (SK4700, Vector Laboratories, USA). After sufficient signal development, the reaction was stopped by rinsing the retinae three times (5 min each) in 0.1M PB, before being mounted in 80% glycerol in 0.1M PB plus 0.1% sodium azide.

## 4.9 Supplementary Tables

**Table S1.** Sampling protocol for assessing the density of neurons in different regions of the retina.

| Cell population | Sampling protocol | No. of retinæ sampled | Counting grid (µm) | Counting frame (µm) | Number of sites sampled | CE<sup>1</sup> |
| --- | --- | --- | --- | --- | --- | --- |
| All photoreceptors | Main | 3 | 800 X 800 | 35 X35 | 230-250 | 0.02-0.03 |
|  | Subsample | 3 | 800 X 800 | 35 X35 | 20-30 | 0.03-0.05 |
| Single cones | Main | 3 | 800 X 800 | 35 X35 | 230-250 | 0.02-0.03 |
|  | Subsample | 3 | 800 X 800 | 35 X35 | 20-30 | 0.03-0.05 |
| Double cones | Main | 3 | 800 X 800 | 35 X35 | 230-250 | 0.03-0.08 |
|  | Subsample | 3 | 800 X 800 | 35 X35 | 20-30 | 0.06-0.09 |
| RH1-expressing photoreceptors | Main | 3 | 850 X 850 | 100 X 100 | 225-250 | 0.03-0.04 |
|  | Subsample | 3 | 300 X 300 | 100 X 100 | 30-35 | 0.03-0.05 |
| SWS1-expressing photoreceptors | Main | 3 | 850 X 850 | 100 X 100 | 190-230 | 0.03-0.04 |
|  | Subsample | 3 | 400 X 400 | 100 X 100 | 12-20 | 0.04-0.09 |

<sup>1</sup> Schaeffer's Coefficient of Error (CE) associated with each sampling protocol (Glaser and Wilson, 1998, Slomianka and West, 2005).

**Table S2.**
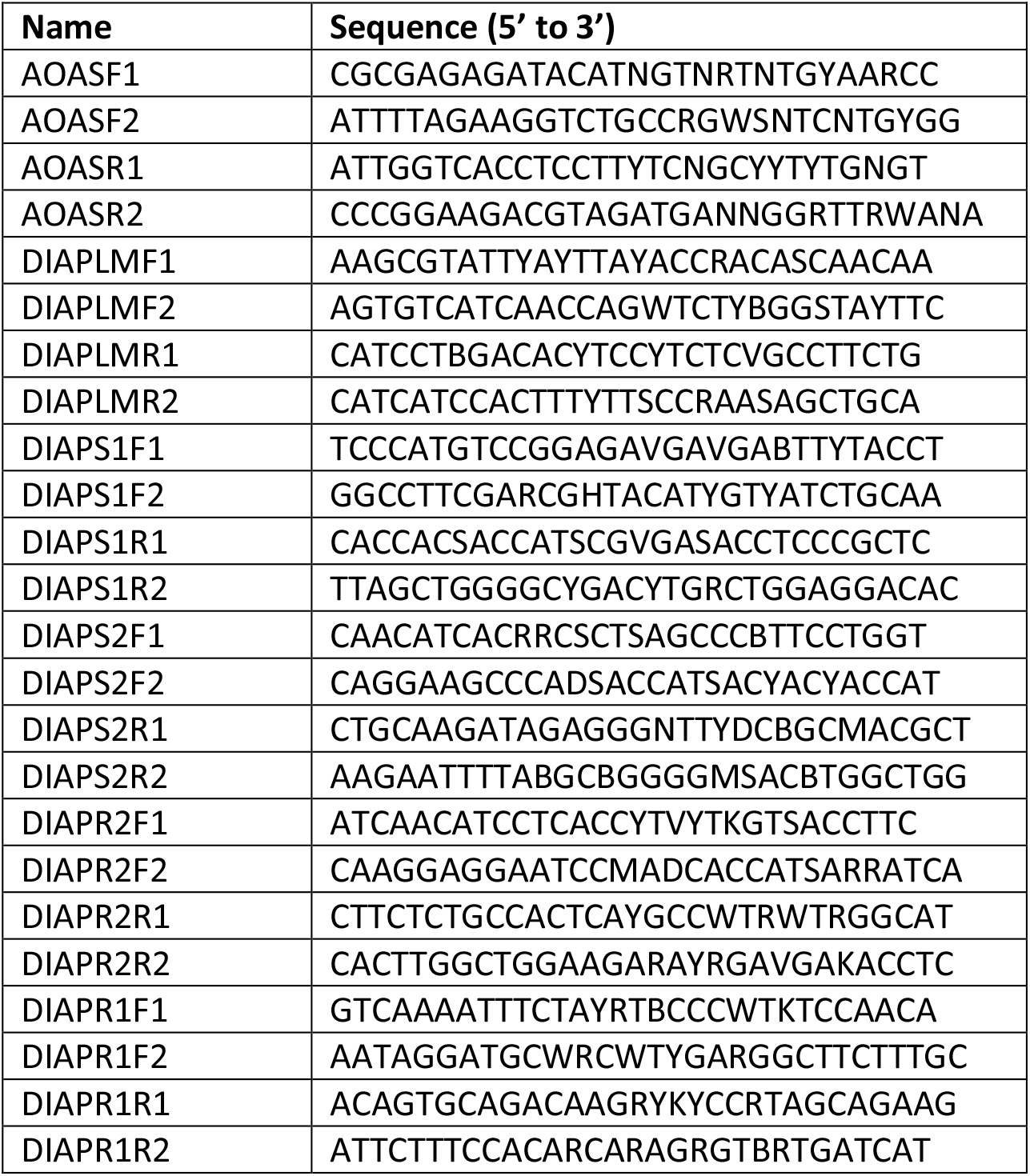
Primers used to isolate and amplify sleepy lizard opsin gene sequences from retinal cDNA (Davies et al., 2009, Knott et al., 2013, Hart et al., 2016).

**Table S3.**
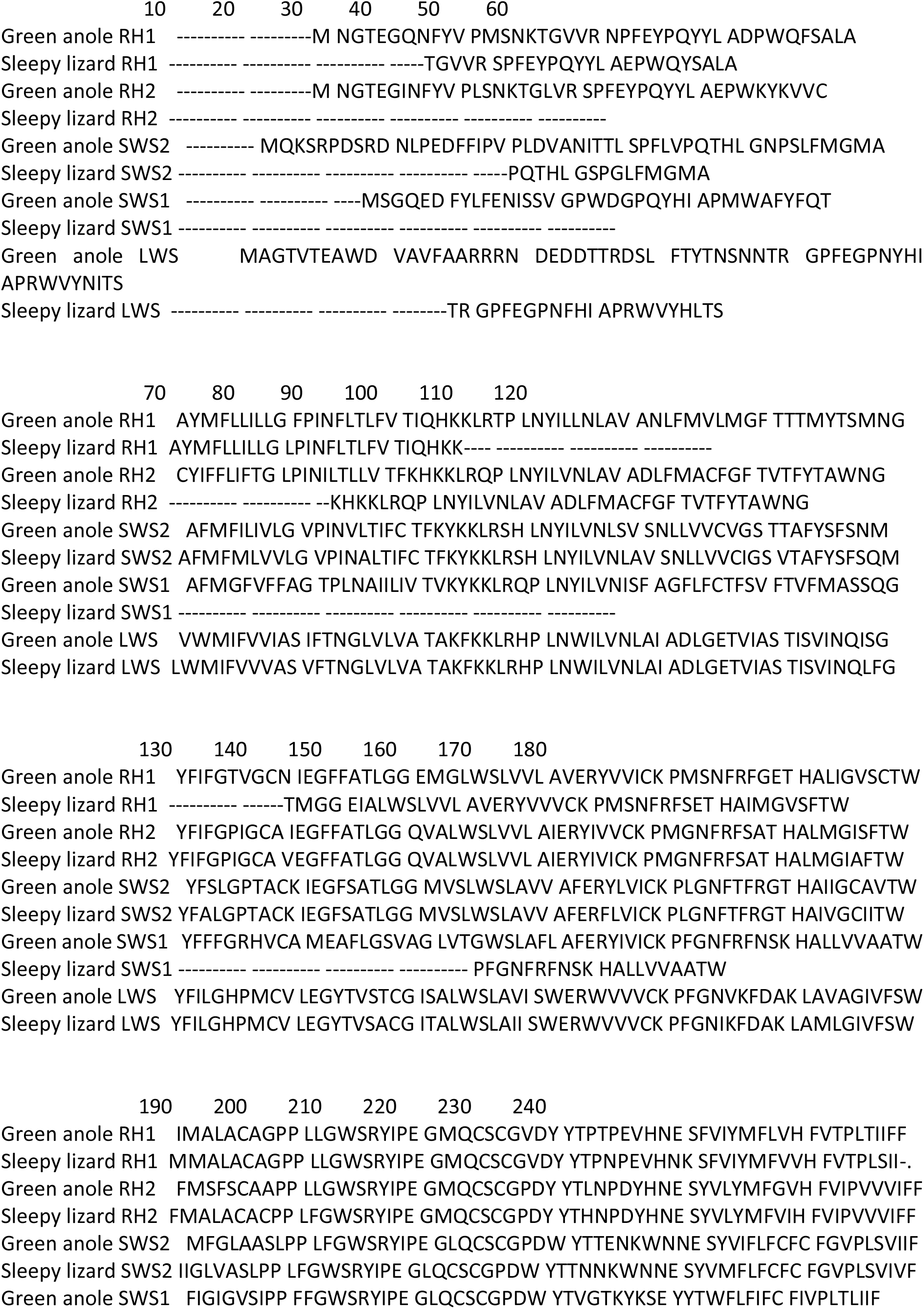
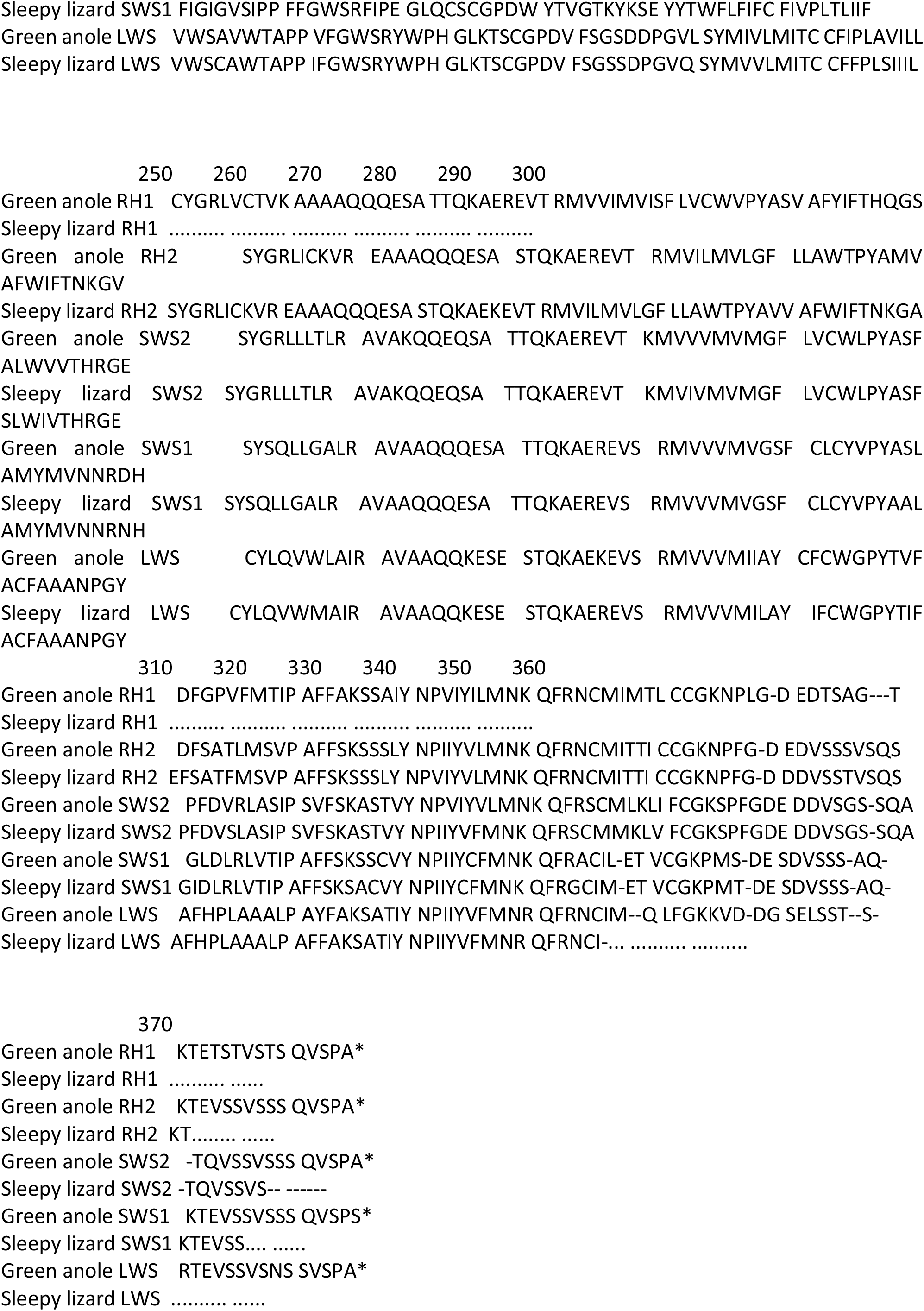
Codon-matched alignment showing partial amino acid sequences for all five visual opsins 908 expressed in the retina of the sleepy lizard, *T. rugosa* compared to orthologues identified in the green anole, *Anolis carolinensis*.

## References

Abramjan, A., Bauerová, A., Somerová, B. and Frynta, D. (2015). Why is the tongue of blue-tongued skinks blue? Reflectance of lingual surface and its consequences for visual perception by conspecifics and predators. The Science of Nature 102, 1–12.

Arden, G. and Tansley, K. (1962). The electroretinogram of a diurnal gecko. The Journal of General Physiology 45, 1145–1161.

Badiane, A., Carazo, P., Price-Rees, S. J., Ferrando-Bernal, M. and Whiting, M. J. (2018). Why blue tongue? A potential UV-based deimatic display in a lizard. Behavioral ecology and sociobiology 72, 104.

Barbour, H. R., Archer, M. A., Hart, N. S., Thomas, N., Dunlop, S. A., Beazley, L. D. and Shand, J. (2002). Retinal characteristics of the ornate dragon lizard, Ctenophorus ornatus. Journal of Comparative Neurology 450, 334–344.

Bennis, M., Molday, R. S., Versaux-Botteri, C., Repérant, J., Jeanny, J.-C. and McDevitt, D. S. (2005). Rhodopsin-like immunoreactivity in the ‘all cone’retina of the chameleon (Chameleo chameleon). Experimental eye research 80, 623–627.

Bowmaker, J. K. (1995). The visual pigments of fish. Progress in retinal and eye research 15, 1–31.

Bowmaker, J. K., Loew, E. R. and Ott, M. (2005). The cone photoreceptors and visual pigments of chameleons. Journal of Comparative Physiology A 191, 925–932.

Coimbra, J. P., Collin, S. P. and Hart, N. S. (2015). Variations in retinal photoreceptor topography and the organization of the rod-free zone reflect behavioral diversity in Australian passerines. Journal of Comparative Neurology 523, 1073–1094.

Coimbra, J. P., Trevia, N., Videira Marceliano, M. L., da Silveira Andrade-Da-Costa, B., Picanço-Diniz, C. W. and Yamada, E. S. (2009). Number and distribution of neurons in the retinal ganglion cell layer in relation to foraging behaviors of tyrant flycatchers. Journal of Comparative Neurology 514, 66–73.

Corbo, J. C. (2021). Vitamin A1/A2 chromophore exchange: Its role in spectral tuning and visual plasticity. Developmental Biology 475, 145–155.

Crescitelli, F. (1972). The Visual Cells and Visual Pigments of the Vertebrate Eye. In Photochemistry of Vision, (ed. H. J. A. Dartnall), pp. 245–363. Berlin, Heidelberg: Springer Berlin Heidelberg.

Crescitelli, F. (1977). The visual pigments of geckos and other vertebrates: An essay in comparative biology. Handbook of sensory physiology 7, 5.

Davies, W. L., Cowing, J. A., Bowmaker, J. K., Carvalho, L. S., Gower, D. J. and Hunt, D. M. (2009). Shedding light on serpent sight: the visual pigments of henophidian snakes. The Journal of Neuroscience 29, 7519–7525.

Ellingson, J., Fleishman, L. and Loew, E. (1995). Visual pigments and spectral sensitivity of the diurnal gecko Gonatodes albogularis. Journal of Comparative Physiology A 177, 559–567.

Fleishman, L., Bowman, M., Saunders, D., Miller, W., Rury, M. and Loew, E. (1997). The visual ecology of Puerto Rican anoline lizards: habitat light and spectral sensitivity. Journal of Comparative Physiology A 181, 446–460.

Fleishman, L. J., Loew, E. R. and Whiting, M. J. (2011). High sensitivity to short wavelengths in a lizard and implications for understanding the evolution of visual systems in lizards. Proceedings of the Royal Society of London B: Biological Sciences 278, 2891–2899.

Forbes, A., Fox, S., Milburn, N. and Deane, H. (1960). Electroretinograms and spectral sensitivities of some diurnal lizards. Journal of neurophysiology 23, 62–73.

Garza-Gisholt, E., Hemmi, J. M., Hart, N. S. and Collin, S. P. (2014). A comparison of spatial analysis methods for the construction of topographic maps of retinal cell density. PLoS One 9, 1–15.

Goldsmith, T. H., Collins, J. S. and Licht, S. (1984). The cone oil droplets of avian retinas. Vision Res 24, 1661–71.

Govardovskii, V., Zueva, L. and Lychakov, D. (1984). Microspectrophotometric study of visual pigments in five species of geckos. Vision Research 24, 1421–1423.

Grötzner, S. R., de Farias Rocha, F. A., Corredor, V. H., Liber, A. M. P., Hamassaki, D. E., Bonci, D. M. O. and Ventura, D. F. (2020). Distribution of rods and cones in the red-eared turtle retina (Trachemys scripta elegans). Journal of Comparative Neurology 528, 1548–1560.

Hamasaki, D. I. (1968). The spectral sensitivity of the lateral eye of the green iguana. Vision Research 8, 1305–1314.

Hárosi, F. I. (1994). An analysis of two spectral properties of vertebrate visual pigments. Vision Research 34, 1359–1367.

Hart, N. S. (2001). Variations in cone photoreceptor abundance and the visual ecology of birds. Journal of Comparative Physiology A 187, 685–697.

Hart, N. S. (2004). Microspectrophotometry of visual pigments and oil droplets in a marine bird, the wedge-tailed shearwater Puffinus pacificus: topographic variations in photoreceptor spectral characteristics. Journal of Experimental Biology 207, 1229–1240.

Hart, N. S., Mountford, J. K., Davies, W. I. L., Collin, S. P. and Hunt, D. M. (2016). Visual pigments in a palaeognath bird, the emu Dromaius novaehollandiae: implications for spectral sensitivity and the origin of ultraviolet vision. Proceedings of the Royal Society B: Biological Sciences 283.

Hauser, F. E., van Hazel, I. and Chang, B. S. W. (2014). Spectral tuning in vertebrate short wavelength-sensitive 1 (SWS1) visual pigments: Can wavelength sensitivity be inferred from sequence data? Journal of Experimental Zoology Part B: Molecular and Developmental Evolution 322, 529–539.

Hauzman, E., Bonci, D. M. O., Suárez-Villota, E. Y., Neitz, M. and Ventura, D. F. (2017). Daily activity patterns influence retinal morphology, signatures of selection, and spectral tuning of opsin genes in colubrid snakes. BMC Evolutionary Biology 17, 249.

Hemmi, J. M. and Grünert, U. (1999). Distribution of photoreceptor types in the retina of a marsupial, the tammar wallaby (Macropus eugenii). Visual neuroscience 16, 291–302.

Hickman, C. P., Roberts, L. S., Keen, S., Larson, A., Helen, I. a. and David, E. (2008). Integrated principles of zoology. USA: McGraw-Hill.

Higgins, D. G., Thompson, J. D. and Gibson, T. J. (1996). Using CLUSTAL for multiple sequence alignments. Methods in Enzymology 266, 383–402.

Howlett, M. H., Smith, R. G. and Kamermans, M. (2017). A novel mechanism of cone photoreceptor adaptation. PloS biology 15, e2001210.

Hunt, D. M. and Peichl, L. E. O. (2014). S cones: Evolution, retinal distribution, development, and spectral sensitivity. Visual neuroscience 31, 115–138.

Jacobs, G. H., Neitz, J. and Krogh, K. (1996). Electroretinogram flicker photometry and its applications. Journal of the Optical Society of America A 13, 641–648.

Jessop, A.-L., Ogawa, Y., Bagheri, Z. M., Partridge, J. C. and Hemmi, J. M. (2020). Photoreceptors and diurnal variation in spectral sensitivity in the fiddler crab Gelasimus dampieri. Journal of Experimental Biology 223.

Katti, C., Stacey-Solis, M., Coronel-Rojas, N. A. and Davies, W. I. L. (2019). The Diversity and Adaptive Evolution of Visual Photopigments in Reptiles. Frontiers in Ecology and Evolution 7.

Kawamura, S. and Yokoyama, S. (1993). Molecular characterization of the red visual pigment gene of the American chameleon (Anolis carolinensis). FEBS letters 323, 247–251.

Kawamura, S. and Yokoyama, S. (1997). Expression of visual and nonvisual opsins in American chameleon. Vision Research 37, 1867–1871.

Kawamura, S. and Yokoyama, S. (1998). Functional characterization of visual and nonvisual pigments of American chameleon (Anolis carolinensis). Vision Res 38, 37–44.

Kirmse, W., Kirmse, R. and Milev, E. (1994). Visuomotor operation in transition from object fixation to prey shooting in chameleons. Biological cybernetics 71, 209–214.

Knott, B., Davies, W. I., Carvalho, L. S., Berg, M. L., Buchanan, K. L., Bowmaker, J. K., Bennett, T. and Hunt, D. M. (2013). How parrots see their colours: novelty in the visual pigments of Platycercus elegans. Journal of Experimental Biology 216, 4454–4461.

Kram, Y. A., Mantey, S. and Corbo, J. C. (2010). Avian Cone Photoreceptors Tile the Retina as Five Independent, Self-Organizing Mosaics. PloS One 5, e8992.

Kryger, Z., Galli-Resta, L., Jacobs, G. and Reese, B. (1998). The topography of rod and cone photoreceptors in the retina of the ground squirrel. Visual neuroscience 15, 685–691.

Loew, E. R., Fleishman, L. J., Foster, R. G. and Provencio, I. (2002). Visual pigments and oil droplets in diurnal lizards a comparative study of Caribbean anoles. Journal of Experimental Biology 205, 927–938.

Martin, M., Le Galliard, J.-F., Meylan, S. and Loew, E. R. (2015). The importance of ultraviolet and near-infrared sensitivity for visual discrimination in two species of lacertid lizards. The Journal of Experimental Biology 218, 458.

McDevitt, D. S., Brahma, S. K., Jeanny, J. C. and Hicks, D. (1993). Presence and foveal enrichment of rod opsin in the “all cone” retina of the American chameleon. The Anatomical Record 237, 299–307.

Mitkus, M., Olsson, P., Toomey, M. B., Corbo, J. C. and Kelber, A. (2017). Specialized photoreceptor composition in the raptor fovea. Journal of Comparative Neurology 525, 2152–2163.

Müller, B. and Peichl, L. (1989). Topography of cones and rods in the tree shrew retina. Journal of Comparative Neurology 282, 581–594.

Murray, K. and Bull, C. M. (2004). Aggressiveness during monogamous pairing in the sleepy lizard, Tiliqua rugosa: a test of the mate guarding hypothesis. Acta ethologica 7, 19–27.

Nei, M. and Kumar, S. (2000). Molecular evolution and phylogenetics: Oxford University Press, USA. 125.

Neitz, J., Geist, T. and Jacobs, G. H. (1989). Color vision in the dog. Visual neuroscience 3, 119-125

New, S. D. and Bull, C. M. (2011). Retinal ganglion cell topography and visual acuity of the sleepy lizard (Tiliqua rugosa). Journal of Comparative Physiology A 197, 703–709.

New, S. T., Hemmi, J. M., Kerr, G. D. and Bull, C. M. (2012). Ocular anatomy and retinal photoreceptors in a skink, the sleepy lizard (Tiliqua rugosa). The Anatomical Record 295, 1727–1735.

Ogawa, Y., Falkowski, M., Narendra, A., Zeil, J. and Hemmi, J. M. (2015). Three spectrally distinct photoreceptors in diurnal and nocturnal Australian ants. In Proc. R. Soc. B, vol. 282, pp. 20150673: The Royal Society.

Osorio, D. (2019). The evolutionary ecology of bird and reptile photoreceptor spectral sensitivities. Current Opinion in Behavioral Sciences 30, 223–227.

Pérez i de Lanuza, G. and Font, E. (2014). Ultraviolet vision in lacertid lizards: evidence from retinal structure, eye transmittance, SWS1 visual pigment genes and _ehavior. The Journal of Experimental Biology 217, 2899.

Provencio, I., Loew, E. R. and Foster, R. G. (1992). Vitamin A 2-based visual pigments in fully terrestrial vertebrates. Vision Research 32, 2201–2208.

Rieke, F. and Baylor, D. A. (2000). Origin and Functional Impact of Dark Noise in Retinal Cones. Neuron 26, 181–186.

Saitou, N. and Nei, M. (1987). The neighbor-joining method: a new method for reconstructing phylogenetic trees. Molecular biology and evolution 4, 406–425.

Schiviz, A. N., Ruf, T., Kuebber-Heiss, A., Schubert, C. and Ahnelt, P. K. (2008). Retinal cone topography of artiodactyl mammals: Influence of body height and habitat. Journal of Comparative Neurology 507, 1336–1350.

Schott, R. K., Müller, J., Yang, C. G., Bhattacharyya, N., Chan, N., Xu, M., Morrow, J. M., Ghenu, A.-H., Loew, E. R. and Tropepe, V. (2016). Evolutionary transformation of rod photoreceptors in the all-cone retina of a diurnal garter snake. Proceedings of the National Academy of Sciences 113, 356–361.

Simões, B. F., Gower, D. J., Rasmussen, A. R., Sarker, M. A. R., Fry, G. C., Casewell, N. R., Harrison, R. A., Hart, N. S., Partridge, J. C., Hunt, D. M. et al. (2020). Spectral Diversification and Trans-Species Allelic Polymorphism during the Land-to-Sea Transition in Snakes. Current Biology 30, 2608-2615.e4.

Skorupski, P. and Chittka, L. (2010). Differences in Photoreceptor Processing Speed for Chromatic and Achromatic Vision in the Bumblebee, <em>Bombus terrestris</em>. The Journal of Neuroscience 30, 3896–3903.

Stapley, J. and Whiting, M. J. (2006). Ultraviolet signals fighting ability in a lizard. Biology Letters 2, 169–172.

Tailby, C., Solomon, S. G. and Lennie, P. (2008). Functional asymmetries in visual pathways carrying S-cone signals in macaque. J Neurosci 28, 4078–87.

Takenaka, N. and Yokoyama, S. (2007). Mechanisms of spectral tuning in the RH2 pigments of Tokay gecko and American chameleon. Gene 399, 26–32.

Tamura, K. and Nei, M. (1993). Estimation of the number of nucleotide substitutions in the control region of mitochondrial DNA in humans and chimpanzees. Molecular biology and evolution 10, 512–526.

Tamura, K., Stecher, G. and Kumar, S. (2021). MEGA11 Molecular Evolutionary Genetics Analysis Version 11. Molecular biology and evolution 38, 3022–3027.

Walls, G. L. (1942). The vertebrate eye and its adaptive radiation. New York: Hafner.

West, M., Slomianka, L. and Gundersen, H. J. G. (1991). Unbiased stereological estimation of the total number of neurons in the subdivisions of the rat hippocampus using the optical fractionator. The Anatomical Record 231, 482–497.

Whitmore, A. and Bowmaker, J. (1989). Seasonal variation in cone sensitivity and short-wave absorbing visual pigments in the rudd Scardinius erythrophthalmus. Journal of Comparative Physiology A 166, 103–115.

Yewers, M. S., McLean, C. A., Moussalli, A., Stuart-Fox, D., Bennett, A. T. D. and Knott, B. (2015). Spectral sensitivity of cone photoreceptors and opsin expression in two colour-divergent lineages of the lizard Ctenophorus decresii. Journal of Experimental Biology 218, 1556–1563.

Zhang, X., Wensel, T. G. and Yuan, C. (2006). Tokay Gecko Photoreceptors Achieve Rod-Like Physiology with Cone-Like Proteins. Photochemistry and photobiology 82, 1452–1460.

